# Subcellular second messenger networks drive distinct repellent-induced axon behaviors

**DOI:** 10.1101/2023.02.02.526245

**Authors:** S Baudet, Y Zagar, F Roche, C Gomez Bravo, S Couvet, J Bécret, M Belle, O Ros, X Nicol

## Abstract

Second messengers, including cAMP, cGMP and Ca^2+^ are often placed in an integrating position to combine the extracellular cues that orient growing axons in the developing brain. This view suggests that axon repellents share the same set of cellular messenger signals and that axon attractants evoke opposite cAMP, cGMP and Ca^2+^ changes. Investigating the confinement of these second messengers in cellular nanodomains, we instead demonstrate that two repellent cues, ephrin-A5 and Slit1, induce spatially segregated signals. These guidance molecules activate subcellular-specific second messenger crosstalks, each signaling network controlling distinct axonal morphology changes *in vitro* and pathfinding decisions *in vivo*.

## Main text

Second messengers, including cyclic nucleotides (cAMP and cGMP) and calcium (Ca^2+^), are key signaling molecules involved in a wide range of cellular pathways. Although diffusing freely in aqueous buffers, the mechanisms enabling them to achieve specificity for their many downstream cellular processes rely on the compartmentation of these signaling molecules^1,2^. The compartmentation of Ca^2+^ has been identified in a range of cell types with a variety of subcellular locations. In developing neurons, Ca^2+^ transients have been imaged in growth cones, at the tip of the extending axons. Slow transients covering the entire growth cone have been imaged, whereas Ca^2+^ elevations of smaller spatial spread have been identified in filopodia and at cell adhesion sites^3–5^. cAMP nanodomains have been described in a variety of forms including biomolecular condensates of high concentration of this second messenger, local compartments with low cAMP amounts or nanodomains containing receptor-specific signaling units^6–8^. In developing neurons, lipid raft-restricted cAMP signals regulate axon pathfinding^9^. The subcellular compartmentation of cGMP has been less investigated but recent studies have identified distinct submembrane domains of this second messenger in cardiomyocytes^10^. However, the functional relevance of subcellular second messenger compartments is still elusive.

Cyclic nucleotides and Ca^2+^ pathways are highly interdependent^11,12^. A subset of cAMP and cGMP synthesizing enzymes are Ca^2+^-regulated^13,14^. The degradation enzymes shared by cyclic nucleotides induce a crosstalk between these signaling molecules^15^. Both the extracellular Ca^2+^ influx and the release of intracellular stores are influenced by the concentration of cAMP and cGMP^16^. Thus, interacting signaling has been identified in many cell types using cell-wide approaches^17–20^. How second messenger compartmentation influences the subcellular interactions between their signaling pathways has been scarcely approached.

In the developing nervous system cAMP, cGMP and Ca^2+^ are key molecules for the establishment of a precisely connected neuronal network. Among other cellular processes occurring in developing neurons, cyclic nucleotides and Ca^2+^ are critical regulators of axonal behavior when the growth cone at the axon distal end faces guidance molecules^21–23^. These cues are expressed in the developing nervous systems and enable axons to follow a genetically-defined path that guide them towards their targets where they connect an appropriate post-synaptic neuron. They influence the orientation of axon outgrowth by repelling them or promoting axon extension. The influence of second messengers in this process has been mostly investigated using pharmacological approaches that do not enable subcellular manipulations. These investigations highlighted that, like in other cell types, cyclic nucleotides and Ca^2+^ interact to regulate axon pathfinding^24,25^. A few morphologically- or biochemically-defined compartments of the axonal growth cone have been identified as key locations for axon guidance. In developing neurons, filopodia-restricted Ca^2+^ and cAMP signals orient axon outgrowth^5,24^ and cAMP signals restricted to the vicinity of lipid rafts are required for ephrinA5-induced retinal axon repulsion^9^. This fraction of the plasma membrane is also required for the impact of other guidance molecules on growing axons, including Semaphorin-3A and Netrin1^26,27^. Overall, these observations led to draw a model in which second messengers are positioned as integrators of all the molecular cues detected by growing axons, and in which a reversal of the cAMP:cGMP ratio is sufficient to convert axon attraction into repulsion^22,25^.

This model suggests that repellent molecules share a common set of second messenger signals, whereas attractants reverse the ratio of cyclic nucleotide concentration. However, not all axon repellents induce the same morphological changes on developing axons, challenging the idea of a common integrative signal based on second messenger overall concentration. For instance, although Slit2 and Semaphorin-3A both repel the axons of dorsal root ganglia neurons, Slit2 induces a rapid elongation of the filopodia before repulsion, whereas axonal growth cones exposed to Semaphorin-3A do not exhibit this striking behavior^28^. This differential behavior suggests that distinct signaling pathways are involved downstream of different axon repellents. We hypothesize that specific second messenger signals (*e*.*g*. restricted to different subcellular domains), control this diversity of axonal responses. *In vivo*, axon repellents from the Slit and ephrin-A families are involved in non-overlapping developmental stages during the pathfinding of retinal ganglion cell (RGC) axons. Whereas Slits are critical for maintaining axons in the optic nerve and tract when retinal axons reach the optic chiasm^29,30^, ephrin-As contribute to terminate retinal axon growth and control the position of their terminal arbors within their main targets in the brain, the superior colliculus (SC) and visual thalamus^31,32^.

We hypothesize that Slits and ephrin-As initiate distinct and subcellularly-confined second messenger signals that are associated with their specific influence on axon behavior. Here, using a genetically-encoded toolset enabling the subcellular monitoring and manipulation of second messengers, we tested this hypothesis in developing RGC axons. Focusing on repellent axon guidance molecules from two distinct families (ephrin-A and Slit), we provide a comprehensive description of second messenger signals and their subcellular interactions in axons facing these cues. We demonstrate that second messenger signaling is confined in a single membrane compartment for each guidance molecule, but differ from one cue to another. These differences correlate with distinct axonal behaviors in response to each cue. Consistently, manipulating second messengers in the subcellular compartment corresponding to either Slit1 or ephrin-A5 induces axon pathfinding defects matching the role of each of these guidance molecules *in vivo*. These observations challenge the theory that the signaling pathways of all guidance molecules are globally integrated by a single set of second messenger modulations.

### Lipid rafts are the seat of ephrin-A5-induced cGMP and Ca^2+^ signals

Since ephrin-A5 leads to a reduction in cAMP concentration restricted to lipid rafts ^9^, we focused on the same cellular compartment to identify a potential domain where changes in the level of cGMP are confined downstream of this axon guidance molecule. We used ^T^hPDE5^VV^, a cGMP-sensitive FRET biosensor, to monitor the concentration of this second messenger in retinal axons *in vitro*. A plasmid encoding ^T^hPDE5^VV^ was electroporated in E14.5 mouse retina. Retinal explants from the electroporated retina were cultured and growing axons were imaged while exposed to ephrin-A5. Since ^T^hPDE5^VV^ is distributed throughout the entire cytosol, it does not enable to identify the source of cGMP. To this aim, the biosensor was co-electroporated with SponGee, a genetically-encoded cGMP scavenger^33^. Variants of SponGee restricted to lipid rafts (Lyn-SponGee), or to the non-raft fraction of the plasma membrane (SponGee-Kras) are available. The subcellular targeting of Lyn-SponGee relies on a N-terminal fusion of a tandem of palmitoylation-myristoylation sites from Lyn Kinase, whereas SponGee-Kras is restricted to the plasma membrane but excluded from lipid rafts by the C-terminal fusion of a CaaX-polylysine motif derived from K-Ras^33^. The expression of the non-targeted variant of SponGee prevents the elevation of the cGMP concentration independently of its subcellular location, whereas Lyn-SponGee and SponGee-Kras enable to prevent the cGMP changes specifically in lipid rafts or in the non-raft plasma membrane, respectively^33^. When co-expressed with ^T^hPDE5^VV^, the variants of SponGee thus enable to localize the subcellular origin of the cGMP signal (Fig. 1a). When expressed alone, ^T^hPDE5^VV^ detects a transient elevation of cGMP shortly after retinal axons are exposed to ephrin-A5. By contrast, this cGMP elevation is not detected after a sham stimulation (Fig. 1b). This ephrin-A5-induced increase in cGMP concentration is prevented by the cytosolic SponGee and by its lipid raft-targeted variant Lyn-SponGee, but not by the lipid raft-excluded SponGee-Kras, demonstrating that ephrin-A5 exposure leads to an elevation of cGMP in the vicinity of lipid rafts (Fig. 1b).

**Figure 1.**
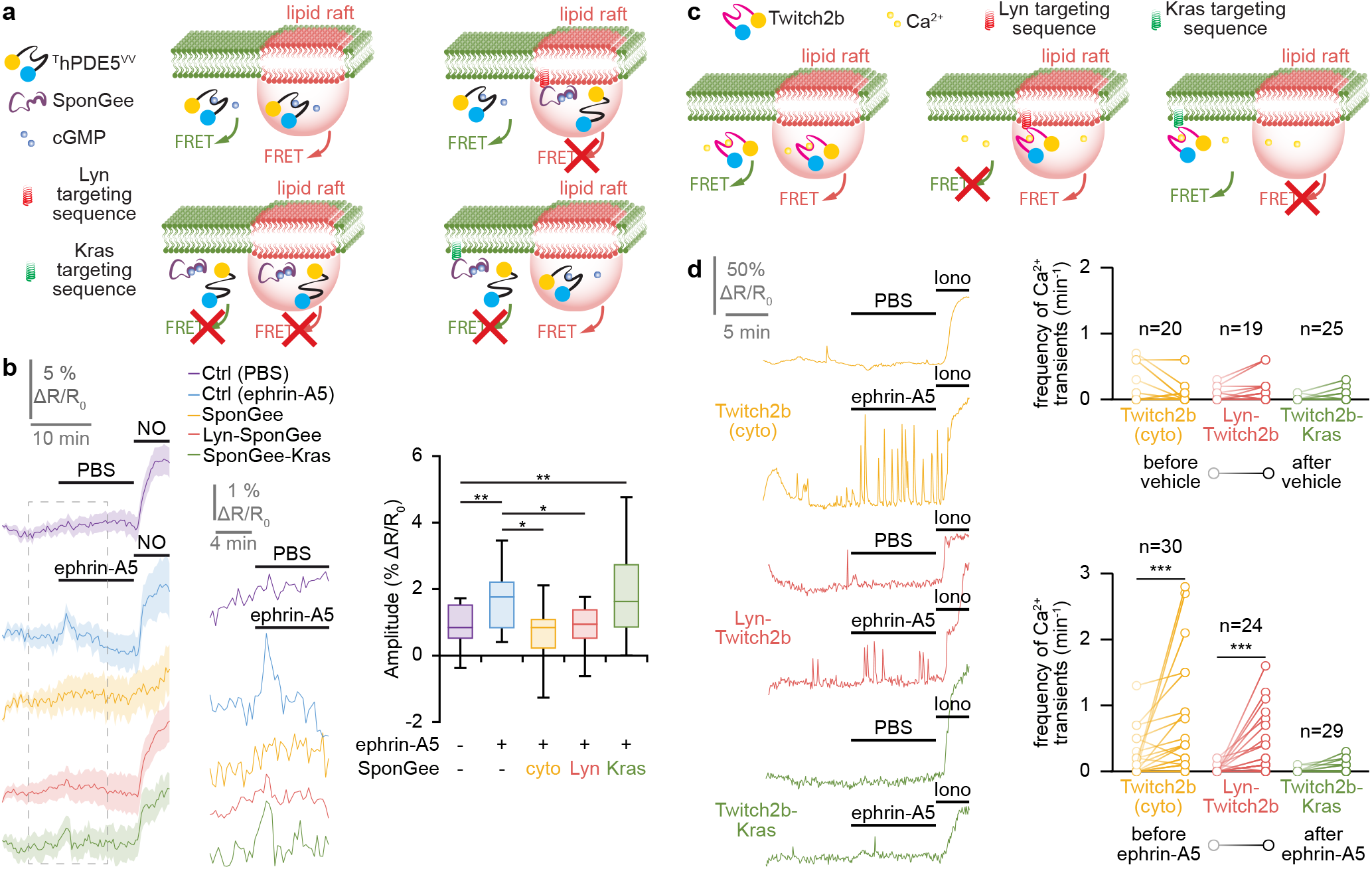
Ephrin-A5 induces an elevation of cGMP and an increase in the frequency of Ca^2+^ transients in lipid rafts. **(a)** Strategy to identify the source of cGMP signals. The cGMP FRET biosensor ^T^hPDE5^VV^ is expressed in RGC axons alone (top left) or with either the non-targeted cGMP scavenger SponGee (bottom left) or its lipid raft-targeted (Lyn-SponGee, top right) or -excluded (SponGee-Kras, bottom right) variants. When expressed alone, ^T^hPDE5^VV^ monitors cGMP changes from the entire cell. When expressed together with SponGee, the sensor is unable to report any change in cGMP concentration. When combined with Lyn-SponGee, ^T^hPDE5^VV^ monitors cGMP changes from the entire cytoplasm excluding the vicinity of lipid rafts. By contrast, when co-expressed with SponGee-Kras, it reflects the fluctuations of cGMP from the cytosol but not from the non-raft fraction of the plasma membrane. **(b)** When retinal axons are exposed to ephrin-A5, ^T^hPDE5^VV^ alone (blue) or co-expressed with SponGee-Kras (green) monitors an elevation of cGMP. By contrast, when co-expressed with SponGee (yellow) or Lyn-SponGee (red), the FRET signal is not affected by ephrin-A5, similarly to axons that are not exposed to ephrin-A5 and express ^T^hPDE5^VV^ (purple). A nitric oxide (NO) stimulation leading to a cGMP elevation is achieved at the end of each recording to verify the functionality of the biosensor and the viability of the axon. The portion of the left traces enclosed in the dashed rectangle is shown magnified in the middle part of the panel. Shadows surrounding traces, s.e.m. Box-and-whisker plot elements: center line, mean; box limits, upper and lower quartiles; whiskers, 10^th^ and 90^th^ percentiles. * *P* < 0.05; ** *P* < 0.01; Kruskal-Wallis test followed by Mann-Whitney post hoc tests. **(c)** Strategy to identify local changes in Ca^2+^ concentration. The Ca^2+^ FRET sensor Twitch2b (left), its lipid raft-targeted (middle) or -excluded (right) variants are expressed in RGCs. They report Ca^2+^ changes from anywhere in the cytosol, from lipid rafts and from the non-raft fraction of the plasma membrane, respectively. **(d)** An elevation in the frequency of Ca^2+^ transients is detected by Twitch2b after ephrin-A5 exposure but not after vehicle (PBS) addition to the culture medium (yellow traces). This observation is reproduced with the lipid raft-targeted Twitch2b (red traces), but not when using its lipid raft-excluded equivalent (green traces). A ionomycin (iono) stimulation leading to a Ca^2+^ elevation is achieved at the end of each recording to verify the functionality of the biosensor and the viability of the axon. Representative traces and individual data points are shown. The number of quantified axons is indicated on the graphs. *** *P* < 0.001; paired Wilcoxon test.

To evaluate whether the same subcellular compartment is also the seat of Ca^2+^ signals, the Ca^2+^ biosensor Twitch2b was used^34^. A lipid raft-restricted variant of Twitch2b (Lyn-Twitch2b) was engineered. The lipid raft targeting of Lyn-Twitch2b was confirmed by membrane fractionation on a sucrose gradient and was found in the same fractions as the lipid raft marker Caveolin (Supplementary Fig. 1). Similarly, Twitch2b was fused to the Kras targeting sequence (Twitch2b-Kras) that shifts the localization of Twitch2b towards the fractions labeled by the non-raft marker Adaptin, demonstrating that this sensor is not targeted to lipid rafts (Supplementary Fig. 1). These two biosensors enable the direct visualization of subcellular Ca^2+^ signals (Fig. 1c), following a strategy recently used with another Ca^2+^ biosensor^35^. Retinal axons expressing either Twitch2b (not targeted) or Lyn-Twitch2b (lipid raft-targeted) exhibited an increase in the frequency of Ca^2+^ transients upon ephrin-A5 exposure (Fig. 1d). These Ca^2+^ transients are characterized by a brief elevation of the Ca^2+^ concentration lasting in the range of 10 seconds and resembles the Ca^2+^ transients previously described in the growth cones of developing axons *in vitro* and *in vivo*^4,24,36^. By contrast, the Kras-targeted variant of Twitch2b did not detect any ephrin-A5-induced change in Ca^2+^ signals (Fig. 1d). Thus, ephrin-A5 induces an increase in the frequency of Ca^2+^ transients that are detected in lipid rafts but not outside of this membrane compartment.

Overall, this set of experiments identifies lipid rafts as a subcellular compartment concentrating the second messenger signals induced by ephrin-As in developing axons. It includes a cAMP reduction^9^, a cGMP elevation and an increase in the frequency of Ca^2+^ transients.

### Slit-induced cAMP, cGMP and Ca^2+^ signals are excluded from lipid rafts

Since the current model of second messenger signaling involved in axon pathfinding places these signaling molecules at the crossroads of many guidance cues, we evaluated whether another retinal axon repellent (Slit1) induces second messenger signals in the same subcellular compartment as ephrin-A5. Using a similar approach as the one described above, we characterized the subcellular features of cAMP, cGMP and Ca^2+^ signals induced by Slit1.

cAMP was monitored using the FRET biosensor H147^37^, for which a lipid raft-targeted (Lyn-H147) and a lipid-raft excluded (H147-Kras) variant are available^9^. Using the cAMP sensor H147 without subcellular targeting, we found that Slit1 induces an overall reduction in cAMP concentration in developing retinal axons (Fig. 2a). This reduction was also detected by a lipid raft-excluded variant of this sensor (H147-Kras), whereas axons expressing the lipid raft-targeted Lyn-H147 exhibited no change in the FRET signal, reflecting a stable cAMP concentration (Fig. 2a). This highlights that Slit1 modulates cAMP in the vicinity of the non-raft fraction of the plasma membrane, but not in the submembrane domain adjacent to lipid rafts, contrasting with the ephrin-A5-induced cAMP modulation^9^.

**Figure 2.**
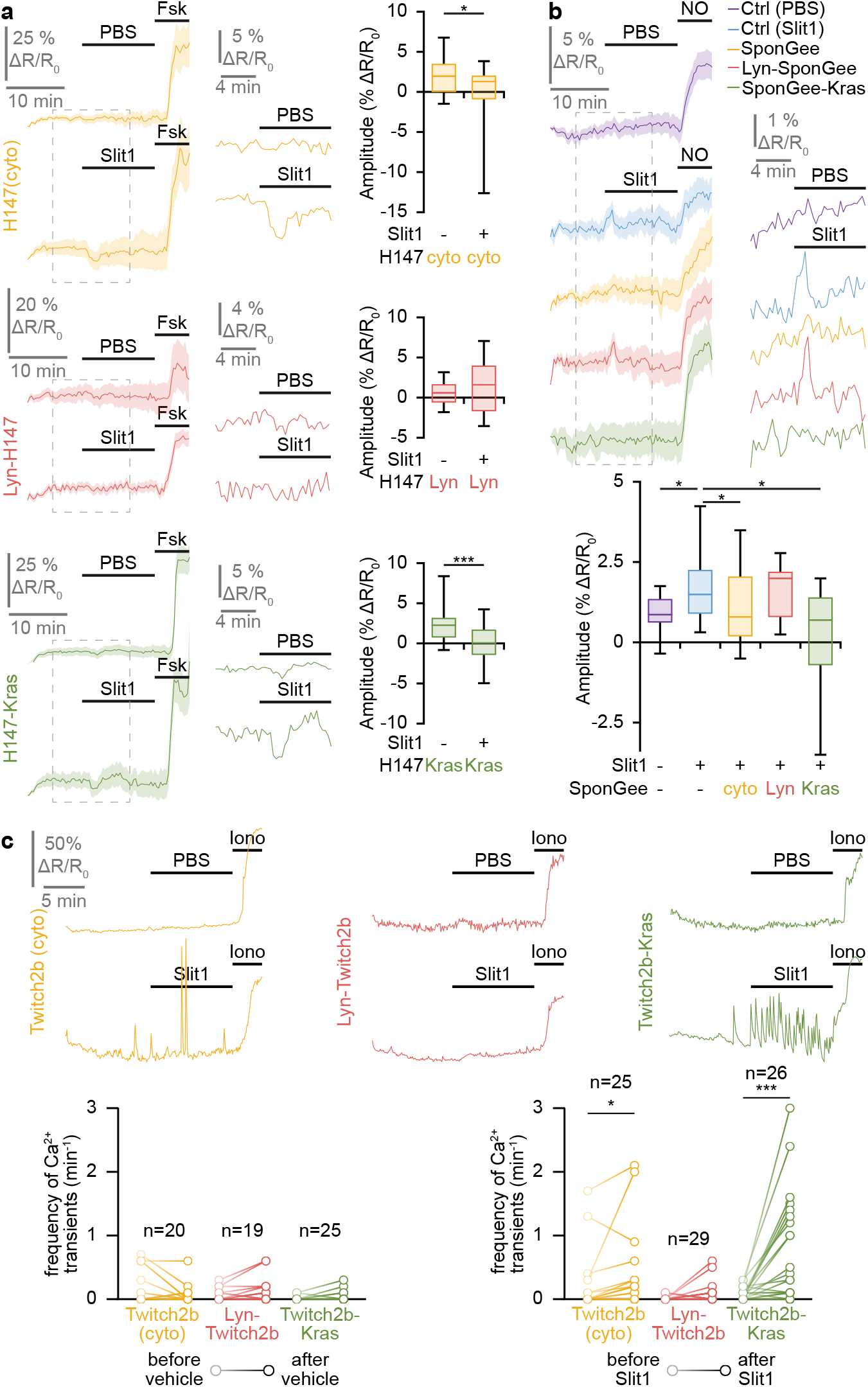
Slit1 induces a cAMP reduction, a cGMP elevation and an increase in the frequency of Ca^2+^ transients in the vicinity of the plasma membrane, outside lipid rafts. **(a)** A reduction in the cAMP concentration is detected by the biosensor H147 after Slit1 exposure but not after vehicle (PBS) addition to the culture medium (yellow traces). This observation is reproduced with the lipid raft-excluded H147 (H147-Kras, green traces), but not when using its lipid raft-targeted equivalent (Lyn-H147, red traces). A forskolin (Fsk) stimulation leading to a cAMP elevation is achieved at the end of each recording to verify the functionality of the biosensor and the viability of the axon. **(b)** When retinal axons are exposed to Slit1, ^T^hPDE5^VV^ alone (blue) or co-expressed with Lyn-SponGee (red) monitors an elevation of cGMP. By contrast, when co-expressed with SponGee (yellow) or Lyn-SponGee (red), the FRET ratio is not affected by Slit1, similarly to axons that are not exposed to Slit1 and express ^T^hPDE5^VV^ (purple). A nitric oxide (NO) stimulation leading to a cGMP elevation is achieved at the end of each recording to verify the functionality of the biosensor and the viability of the axon. **(c)** An elevation in the frequency of Ca^2+^ transients is detected by Twitch2b after Slit1 exposure but not after vehicle (PBS) addition to the culture medium (yellow traces). This observation is reproduced with the lipid raft-excluded Twitch2b (green traces), but not when using its lipid raft-excluded equivalent (red traces). A ionomycin (iono) stimulation leading to a Ca^2+^ elevation is achieved at the end of each recording to verify the functionality of the biosensor and the viability of the axon. **(a**,**b)** The portion of the left traces enclosed in the dashed rectangles is shown magnified in the right part of the panel. Shadows surrounding traces, s.e.m. Box-and-whisker plot elements: center line, mean; box limits, upper and lower quartiles; whiskers, 10^th^ and 90^th^ percentiles. **(c)** Representative traces and individual data points are shown. The number of quantified axons is indicated on the graphs. * *P* < 0.05; *** *P* < 0.001, **(a)** Mann-Whitney test, **(b)** Kruskal-Wallis test followed by Mann-Whitney post hoc tests, **(c)** paired Wilcoxon test.

cGMP concentration was monitored in Slit1-stimulated retinal axons using ^T^hPDE5^VV^. An elevation in cGMP was detected upon Slit1 exposure, mimicking the ephrin-A5-dependent signals (Fig. 2b). However, Slit1-induced cGMP increase was prevented by the raft-excluded scavenger SponGee-Kras, whereas the raft-targeted equivalent Lyn-SponGee was unable to reduce this cGMP elevation (Fig. 2b). This experiment demonstrates that Slit1 controls the cGMP concentration in the vicinity of the non-raft plasma membrane, in contrast to ephrin-A5.

Ca^2+^ concentration was imaged in Slit1-exposed retinal growth cones. Similar to ephrin-A5, an elevation of the frequency of Ca^2+^ transient was detected when using the non-targeted biosensor Twitch2b (Fig. 2c). In contrast to ephrin-A5, the Twitch2b variant targeted in the vicinity of the non-raft plasma membrane (Twitch2b-Kras), but not its Lyn-targeted equivalent, also detected this Ca^2+^ signal, demonstrating that, like for cyclic nucleotides, Slit1-induced Ca^2+^ signals are excluded from lipid rafts (Fig. 2c).

Overall, Slit1 and ephrin-A5, although both exhibiting a repellent activity on developing retinal axons, generate a set of second messenger signals that are restricted to distinct subcellular compartments.

### Subcellular interactions between second messenger signals in developing retinal axons

To further characterize the differences in second messenger signaling downstream of Slit1 and ephrin-A5, we investigated the crosstalks between cyclic nucleotides and Ca^2+^ with subcellular resolution. To this aim, we benefited from a set of genetically-encoded scavengers targeting cyclic nucleotides and Ca^2+^. cAMP Sponge enables to buffer cAMP in living cells^38^ and is available as lipid raft-targeted (Lyn-cAMP Sponge) and -excluded (cAMP Sponge-Kras) variants^9^. SponGee and its variants enable the subcellular manipulation of cGMP^33^. SpiCee is a Ca^2+^ scavenger that has been fused to the lipid raft-targeted and -excluded sequences (Lyn-SpiCee and SpiCee-Kras, respectively)^39^. The ability of the targeted scavengers to prevent second messenger signals was verified by the co-expression of a given scavenger with the corresponding biosensors: ^T^hPDE5^VV^ for cGMP (Fig. 1b; Fig. 2b) or the targeted variants of H147 and Twitch2b for cAMP and Ca^2+^ respectively (Supplementary Fig. 2).

To evaluate the interactions between the second messenger signals, RGCs expressing a subcellular-specific biosensor sensitive to second messenger A were co-electroporated with a scavenger preventing the downstream signaling of second messenger B and exposed to either Slit1 or ephrin-A5. For instance Lyn-Twitch2b was co-expressed with Lyn-cAMP Sponge in axons exposed to ephrin-A5 to determine whether lipid raft-specific cAMP signaling influences the ephrin-A5-induced elevation in the frequency of Ca^2+^ transients in the same subcellular compartment.

Since all second messenger signals detected downstream of ephrin-A5 were found in lipid rafts, we investigated the crosstalk between these signaling molecules within this cellular compartment. Buffering cAMP with Lyn-cAMP sponge was sufficient to inhibit both the elevation of cGMP detected by ^T^hPDE5^VV^ and the increase in Ca^2+^ transient frequency monitored by Lyn-Twitch2b (Fig. 3a,b). By contrast, scavenging cGMP or Ca^2+^ in lipid rafts with Lyn-SponGee or Lyn-SpiCee did not affect the ephrin-A5-induced signals for either of the second messengers (cGMP, cAMP and Ca^2+^) (Fig. 3a-c). Thus, in lipid rafts of retinal growth cones exposed to ephrin-A5, cAMP is positioned upstream of cGMP and Ca^2+^ (Fig. 3d).

**Figure 3.**
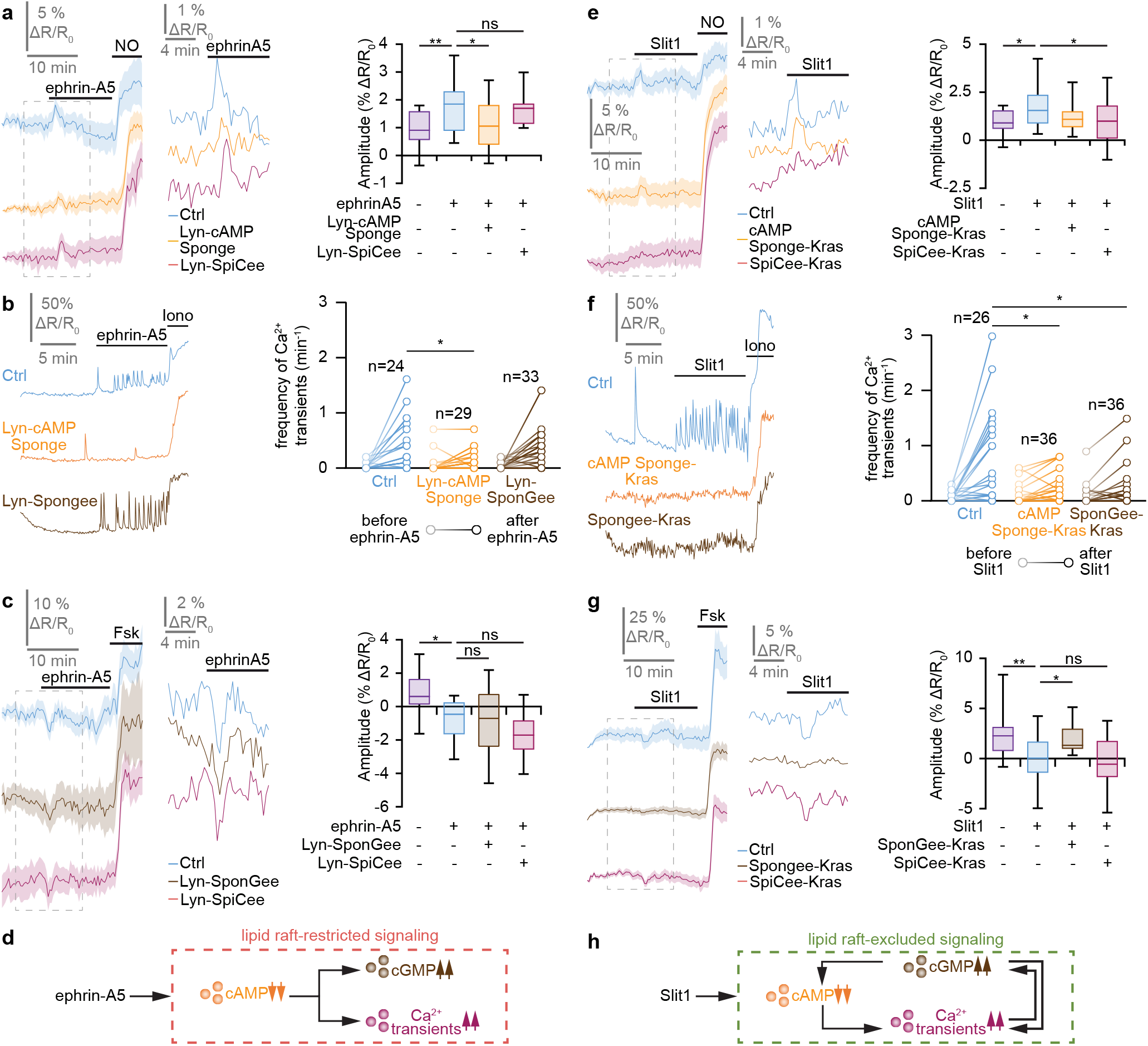
Lipid raft-restricted and -excluded second messenger networks downstream of ephrin-A5 and Slit1. **(a)** The overall cGMP changes induced by ephrin-A5 were monitored using ^T^hDPE5^VV^. The elevation of the cGMP concentration was reduced by preventing local cAMP signals in lipid rafts using Lyn-cAMP Sponge. By contrast, reducing Ca^2+^ downstream signaling in this cellular domain does not affect the cGMP changes. A nitric oxide (NO) stimulation leading to a cGMP elevation is achieved at the end of each recording to verify the functionality of the biosensor and the viability of the axon. **(b)** The Ca^2+^ signals induced by ephrin-A5 in lipid rafts were recorded using Lyn-Twitch2b. The elevation in the Ca^2+^ transient frequency was prevented by scavenging cAMP in lipid rafts using Lyn-cAMP Sponge, whereas altering the downstream signaling of cGMP in this subcellular domain did not impact the ephrin-A5-induced Ca^2+^ transients. A ionomycin (iono) stimulation leading to a Ca^2+^ elevation is achieved at the end of each recording to verify the functionality of the biosensor and the viability of the axon. **(c)** The lipid raft-restricted cAMP signals induced by ephrin-A5 were monitored using the biosensor Lyn-H147. The reduction in the cAMP concentration was not affected by preventing the cGMP or Ca^2+^ downstream signaling in the same cellular compartment using Lyn-SponGee and Lyn-SpiCee, respectively. A forskolin (Fsk) stimulation leading to a cAMP elevation is achieved at the end of each recording to verify the functionality of the biosensor and the viability of the axon. **(d)** Overall model of the lipid raft-restricted second messenger network downstream of ephrin-A5. Exposing growth cones to this axon guidance molecule leads to a combined modulation of cyclic nucleotides and Ca^2+^ that is restricted to the vicinity of lipid rafts. This network is characterized by a non-reciprocal influence of cAMP on cGMP and Ca^2+^ signaling. **(e)** The Slit1-dependent changes in cGMP were imaged using ^T^hDPE5^VV^. The elevation in cGMP was prevented by the blockade of Ca^2+^ next to the non-raft domain of the plasma membrane using SpiCee-Kas. By contrast, blocking cAMP downstream signaling outside lipid rafts with cAMP Sponge-Kras did not affect the Slit1-induced cGMP changes. A nitric oxide (NO) stimulation leading to a cGMP elevation is achieved at the end of each recording to verify the functionality of the biosensor and the viability of the axon. **(f)** The Ca^2+^ signals induced by Slit1 outside lipid rafts were recorded using Twitch2b-Kras. The elevation in the Ca^2+^ transient frequency was prevented by scavenging either cAMP in lipid rafts using Lyn-cAMP Sponge or cGMP using SponGee-Kras. A ionomycin (iono) stimulation leading to a Ca^2+^ elevation is achieved at the end of each recording to verify the functionality of the biosensor and the viability of the axon. **(g)** The juxta-membrane lipid raft-excluded cAMP signals induced by Slit1 were monitored using the biosensor H147-Kras. The reduction in the cAMP concentration was reduced by preventing cGMP downstream signaling in the same cellular compartment using SponGee-Kras, but not by reducing Ca^2+^ signaling with SpiCee-Kras. A forskolin (Fsk) stimulation leading to a cAMP elevation is achieved at the end of each recording to verify the functionality of the biosensor and the viability of the axon. **(h)** Overall model of the juxta-membrane lipid raft-excluded second messenger network downstream of Slit1. Exposing growth cones to this axon guidance molecule leads to a combined modulation of cyclic nucleotides and Ca^2+^ that is restricted to the vicinity of the non-raft domain of the plasma membrane. This network is characterized by complex interactions including a cAMP influence on Ca^2+^, a control of cGMP elevation by Ca^2+^ transients and cGMP influencing both cAMP and Ca^2+^ signals. **(a**,**c**,**e**,**g)** The portion of the left traces enclosed in the dashed rectangle in is shown magnified in the right part of the panel. Shadows surrounding traces, s.e.m. Box-and-whisker plot elements: center line, mean; box limits, upper and lower quartiles; whiskers, 10^th^ and 90^th^ percentiles. **(b**,**f)** Representative traces and individual data points are shown. The number of quantified axons is indicated on the graphs. **(a-c, b-g)** * *P* < 0.05; ** *P* < 0.01; *** *P* < 0.001; Kruskal-Wallis test followed by Mann-Whitney post hoc tests.

In contrast to ephrin-A5, Slit1 modulates cyclic nucleotides and Ca^2+^ in the non-raft fraction of the plasma membrane. Accordingly, ^T^hPDE5^VV^, Twitch2b-Kras and H147-Kras were used to evaluate the crosstalk between Slit1-induced second messenger signals. Buffering cAMP changes in the non-raft plasma membrane with cAMP-Sponge-Kras did not affect the increase in cGMP but reduced the elevation of the Ca^2+^ transient frequency induced by Slit1 (Fig. 3e,f). Scavenging cGMP outside lipid rafts with SponGee-Kras prevented both the Ca^2+^ and cAMP changes in retinal axons (Fig. 3f,g). Preventing Ca^2+^ signaling in the non-raft membrane by expressing SpiCee-Kras precluded the elevation of cGMP concentration but did not affect the reduction in cAMP (Fig. 3e,g). Thus, Slit1 induces the activation of a complex signaling network intermingling cyclic nucleotides and Ca^2+^ signaling. The relationships between the signaling molecules involved in this network are illustrated in Figure 3h.

Overall, we demonstrate that not only are second messenger modulations downstream of ephrin-A5 and Slit1 confined to distinct subcellular compartments, but also that the signaling networks formed by these molecules differ downstream of each of these axon guidance cues.

### Domain-specific signals are required for axon repulsion

To determine whether the lipid raft-restricted and -excluded second messenger signals are required for ephrin-A5- and Slit1-induced retraction, the behavior of retinal axons exposed to these guidance molecules was evaluated. RGC axons were grown *in vitro* after electroporation of the retina with either mRFP or one of the cytosolic or subcellular compartment-targeted scavengers. When exposed to repellents *in vitro*, axonal growth cones undergo a morphological change that is characterized by the loss of their lamellipodium. The ephrin-A5- or Slit1-induced growth cone collapse was evaluated 20 minutes after RGC axon exposure to the guidance cue, *i*.*e*. shortly after the detected second messenger signals. SpiCee, SponGee and cAMP sponge or their Lyn- or Kras-targeted variants were expressed in retinal axons exposed to guidance molecules. None of these second messenger buffers affects the morphology of retinal growth cones that were not exposed to Slit1 or ephrin-A5 (Supplementary Fig. 3a).

Ephrin-A5 induces the collapse of mRFP-expressing growth cones. The fraction of collapsed axons was reduced when SpiCee, SponGee or their lipid raft-restricted variants were expressed (Fig. 4a,b). By contrast, SpiCee-Kras and SponGee-Kras did not affect the response of retinal axons to ephrin-A5. Similarly, Lyn-cAMP sponge, but not cAMP sponge-Kras, was previously reported to prevent the retraction of retinal axons induced by ephrin-A5^9^. The binding of Ca^2+^ and cGMP to Lyn-SpiCee and Lyn-SponGee, respectively, is critical to preclude the collapse of RGC growth cones since this change in the axon morphology was unaffected in axons expressing variants of these scavengers carrying point mutations abolishing their ability to bind their targets (Lyn-mut SpiCee and Lyn-mut SponGee, Supplementary Fig. 3b). Overall, this set of experiments demonstrates that the lipid raft-restricted cyclic nucleotide and Ca^2+^ signals detected after exposure to ephrin-A5 are required for axon repulsion.

**Figure 4.**
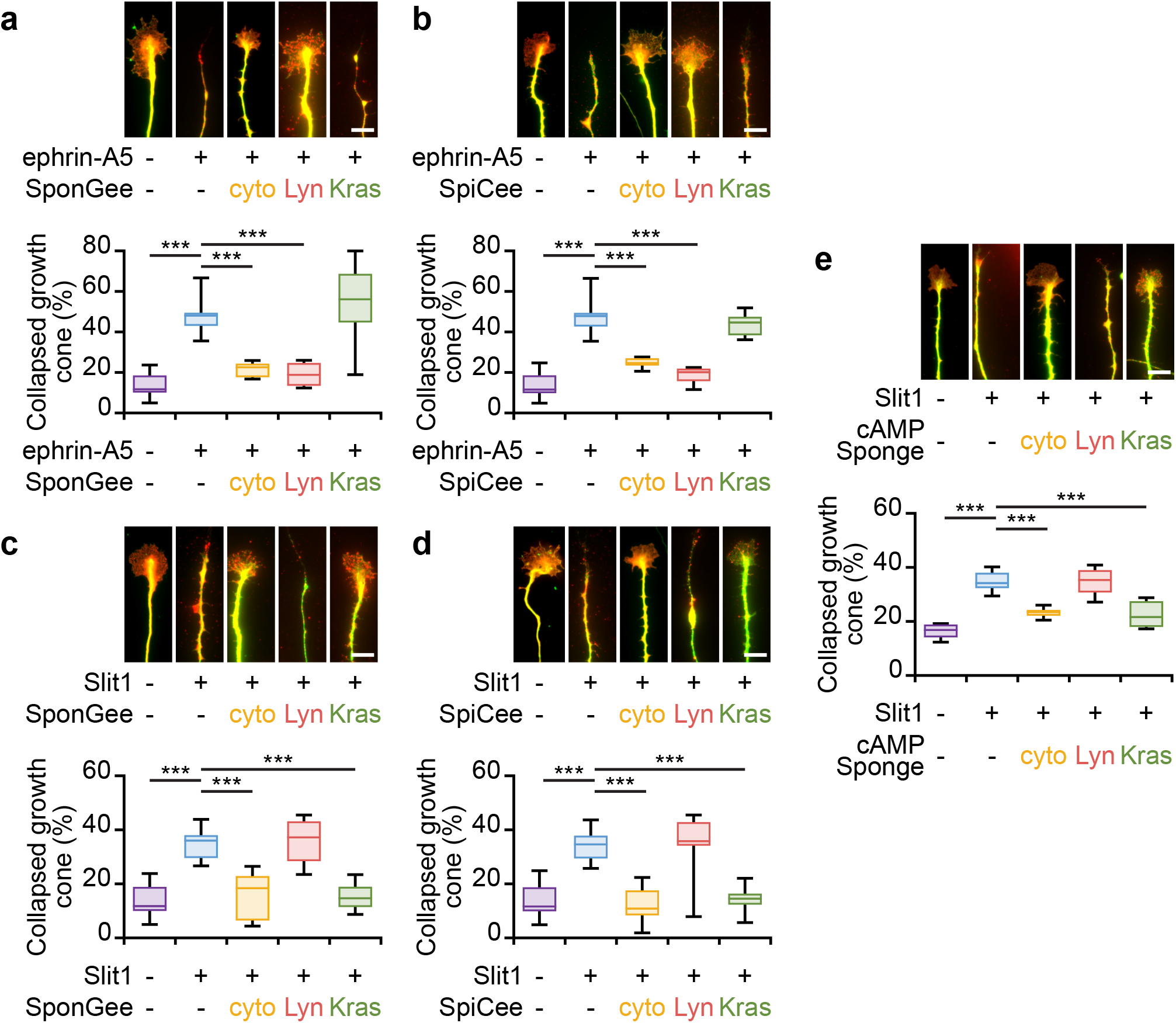
Lipid raft-specific and -excluded scavenging of second messengers prevent the collapse of growth cones induced by ephrin-A5 and Slit-1, respectively. **(a)** Ephrin-A5 induces the collapse of mRFP- and SponGee-Kras-expressing axons, whereas the non-targeted and lipid-raft targeted variants of SponGee (Lyn-SponGee) prevent growth cone collapse. **(b)** When lacking a targeting sequence or when restricted to lipid rafts, SpiCee prevents the ephrin-A5-induced growth cone collapse, in contrast to the lipid raft-excluded variant of SpiCee (SpiCee-Kras). **(c)** Slit-1 induces the collapse of mRFP-expressing retinal growth cones. The proportion of collapsing axons is not affected by the expression of Lyn-SponGee but is reduced by SponGee-Kras or the cytosolic SponGee. **(d)** The collapse of retinal growth cones exposed to Slit-1 is reduced by the expression of SpiCee (not targeted) or its lipid raft-excluded variant (SpiCee-Kras), but not by Lyn-SpiCee. **(e)** Slit-1-induced growth cone collapse is prevented by cAMP Sponge or cAMP Sponge-Kras, but not by Lyn-cAMP-Sponge. Axons were immunolabeled with a βIII-tubulin (green) and a Ds-Red (red) antibody. The latter reports the expression of SponGee **(a**,**c)**, SpiCee **(b**,**d)** or cAMP Sponge **(e)**. Scale bars, 10 μm. Box-and-whisker plot elements: center line, mean; box limits, upper and lower quartiles; whiskers, 10^th^ and 90^th^ percentiles. *** *P* < 0.001; One way ANOVA.

When exposed to Slit1 for 20 minutes, mRFP-expressing axons collapse, whereas a higher number of axons with an intact growth cone were observed when SpiCee, SponGee, cAMP sponge or their variant targeted to the non-raft plasma membrane are expressed (Fig. 4c–e). By contrast, Lyn-SpiCee, Lyn-SponGee and Lyn-cAMP sponge are not able to prevent the Slit1-induced growth cone collapse of retinal axons (Fig. 4c–e), demonstrating that the second messenger signals required for the Slit1-induced collapse of retinal axons are restricted to the non-raft domain of the plasma membrane. This conclusion is confirmed by the unaltered collapse of growth cones expressing variants of SpiCee-Kras, SponGee-Kras and cAMP Sponge-Kras that carry point mutations preventing their ability to bind Ca^2+^, cGMP and cAMP, respectively (mut SpiCee-Kras, mut SponGee-Kras and mut cAMP Sponge-Kras, Supplementary Fig. 3b)

Altogether, we here show that the cGMP, cAMP and Ca^2+^ signals inducing the collapse of the growth cone are confined to lipid rafts when the retraction is driven by ephrin-A5, whereas the compartment involved is the non-raft fraction of the plasma membrane downstream of Slit1.

### Slit1 and ephrin-A5 induce distinct morphological changes of RGC growth cones

The collapse assay provides a coarse characterization of axonal repulsion. Since ephrin-A5- and Slit1-induced growth cone collapse relies on second messenger signaling in distinct cellular compartments, these axon guidance molecules might induce axon repulsion with distinct features. To evaluate whether the response of retinal growth cones exposed to ephrin-A5 diverges from the effect of an exposure to Slit1, the behavior of living axons facing these guidance molecules was monitored during 20 minutes after adding the axon repellent to the culture medium. Control axons growing in a medium supplemented with PBS continue to extend. By contrast, ephrin-A5 exposed axons quickly collapse and retract, with a fast backward movement (Fig. 5; Supplementary Movie 1). Axons exposed to Slit1 exhibit a different behavior: when facing Slit1, they stop growing and collapse with little or no retraction in the 20 minutes following the addition of Slit1 to the culture medium (Fig. 5; Supplementary Movie 1). This differential behavior matches the role of ephrin-A5 and Slit1 during the development of retinal axons. Slit1 is involved in orienting the growth of the axons and in preventing them to exit the corridor formed by the optic nerve, chiasm and tract, but does induce the retraction of retinal axons that have not yet reached their targets. By contrast, ephrin-A5 is expressed in regions of the brain where retinal axons are strongly repelled and exhibit the retraction of branches overshooting the position of their mature terminal arbor^40^.

**Figure 5.**
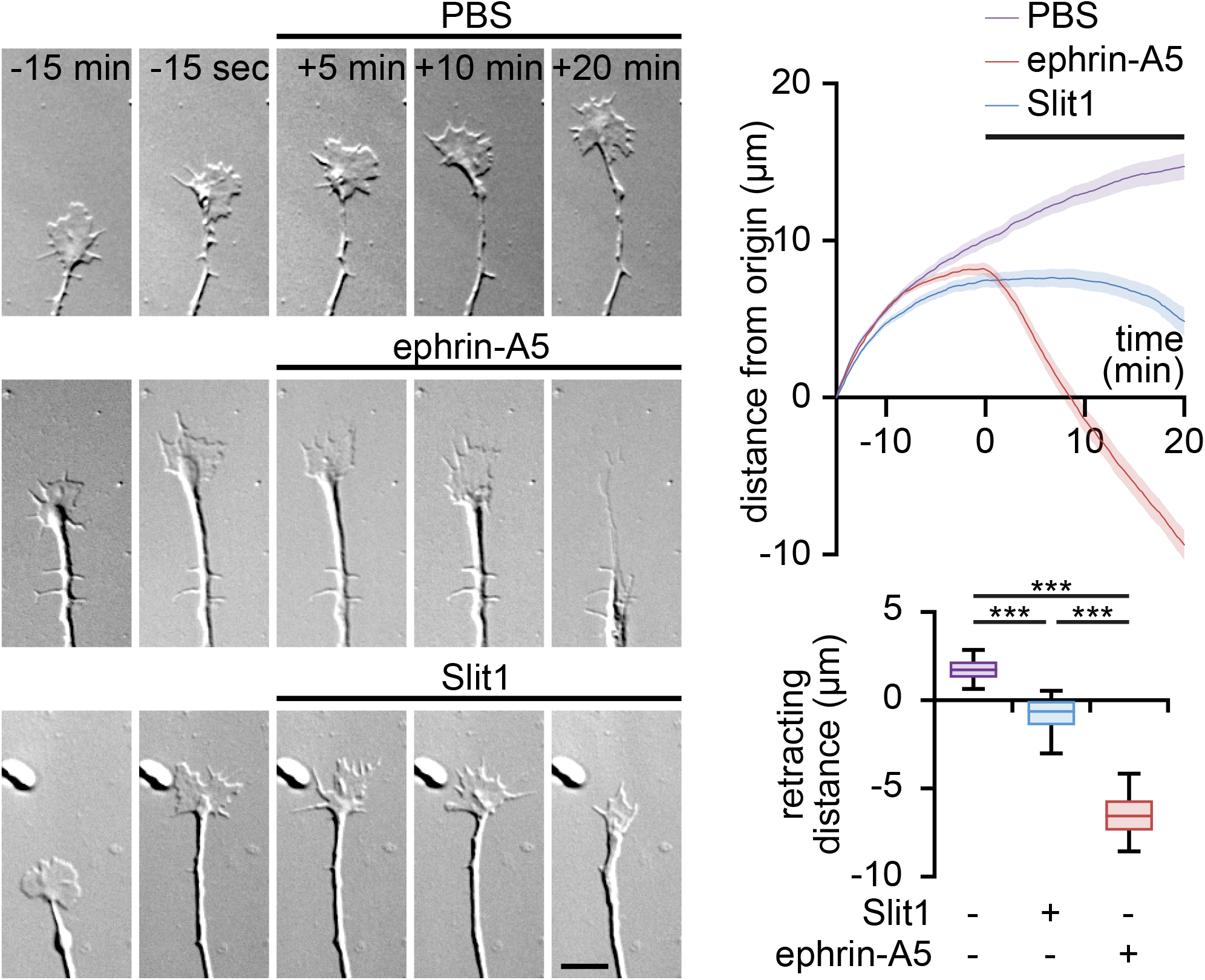
Ephrin-A5 and Slit1 induce distinct morphological changes of axonal growth cones *in vitro*. The growth of axons exposed to PBS was not affected (top row, purple trace). Ephrin-A5 induced a growth cone collapse followed by a prompt retraction (middle row, red trace). Axons exposed to Slit1 exhibited a collapse of the growth cones but in contrast to axons encountering ephrin-A5, did not retract within the 20 minutes recorded. Shadows surrounding traces, s.e.m. Box-and-whisker plot elements: center line, mean; box limits, upper and lower quartiles; whiskers, 10^th^ and 90^th^ percentiles. *** *P* < 0.001; Kruskal-Wallis test followed by Mann-Whitney post hoc tests.

### Impact of second messenger signaling on the development of retinal axon arbors in vivo

To evaluate whether the second messenger signals confined to or excluded from lipid rafts regulate distinct repulsion behaviors *in vivo*, the lipid raft-targeted or -excluded version of SponGee or SpiCee was electroporated *in utero* in the developing retina of E14.5 embryos. Regions of the brain where retinal axon pathfinding relies on Slit1 or ephrin-A5 were analyzed after whole brain clearing and light-sheet imaging: the SC where ephrin-As are critical to shape retinotopic mapping and prevent axons from invading the inferior colliculus, and the optic chiasm where retinal axons are confined within their correct path by Slit1 and Slit2. mRFP-electroporated axons arborize in the SC and form dense terminal arbors at P15. No axonal branches were found in the inferior colliculus, where ephrin-As are highly expressed. By contrast, retinal axons expressing either Lyn-SponGee or Lyn-SpiCee exhibited exuberant branches in the inferior colliculus. In a few cases with sparse electroporation, multiple termination zones for a single axon were found in the SC. These abnormal retinal axon projections were not detected in animals electroporated with the raft-excluded SponGee-Kras and SpiCee-Kras (Fig. 6a). Thus, preventing second messenger signaling specifically in lipid rafts induces a phenotype matching the alterations of the retinal projections in animals lacking a subset of ephrin-A receptors^41^.

**Figure 6.**
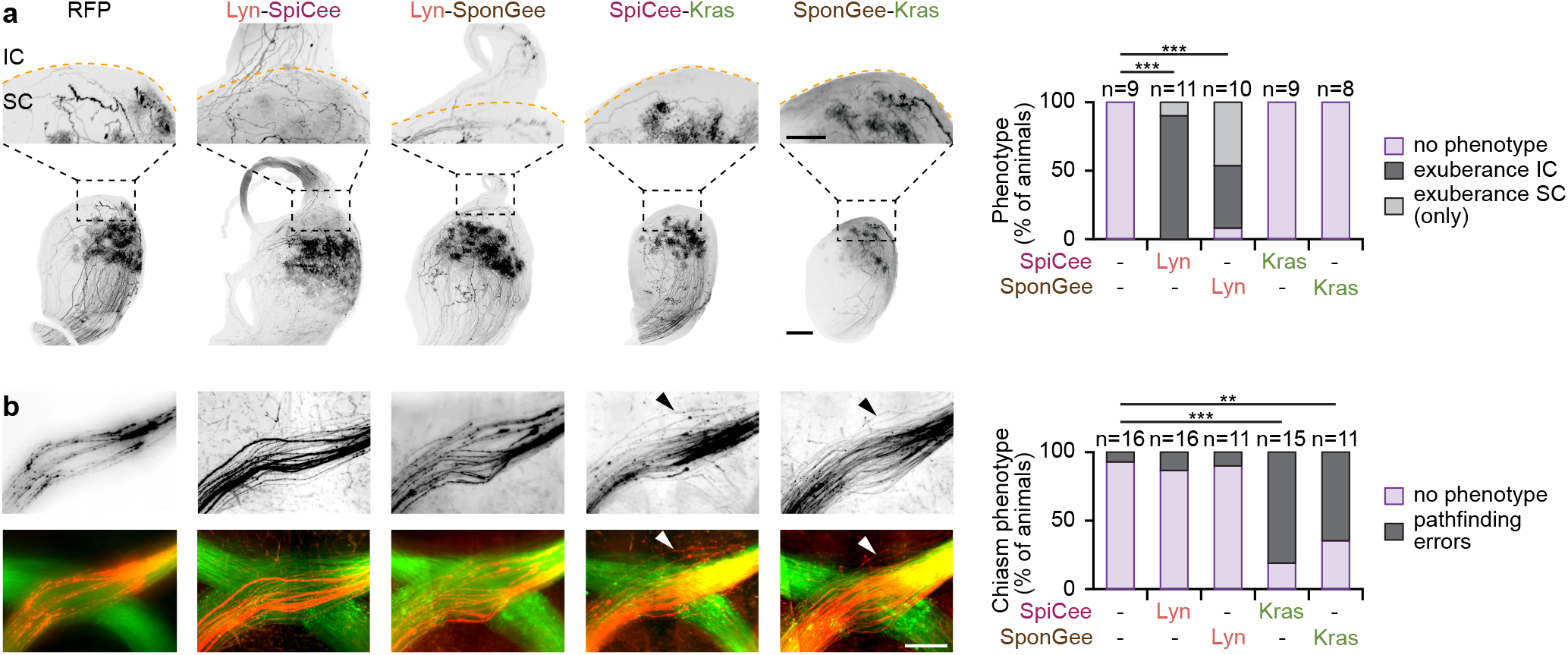
Lipid raft-targeted and -excluded scavenging lead to misguided retinal axons in the SC and at the optic chiasm, respectively. **(a)** Lyn-SpiCee- and Lyn-SponGee-expression lead to overshooting axons in the inferior colliculus at P15, by contrast to mRFP-, SpiCee-Kras- and SponGee-Kras-expression. The orange dashed line highlights the position of the posterior end of the superior colliculus (SC). The inferior colliculus (IC) is above this line. The top row images are magnifications of the regions of the bottom row images indicated by the black dashed squares. Scale bars: top row, 250 μm; bottom row, 500 μm. **(b)** SpiCee-Kras- and SponGee-Kras-expressing axons (red) exit the optic chiasm labeled with TAG1 (green), by contrast to the axons of mRFP-, SpiCee-Kras- and SponGee-Kras-electroporated RGCs. The top row highlights the mRFP channels in which electroporated axons are seen. Scale bar, 200 μm. **(a**,**b)** ** *P* < 0.01; *** *P* < 0.001; χ^2^ test followed by χ^2^ post hoc tests.

The chiasm of animals expressing the lipid raft-targeted or -excluded variant of SponGee or SpiCee were imaged at E18.5 using light sheet microscopy after whole brain clearing. To visualize the position of the unaffected chiasm, a TAG1 immunostaining was performed, enabling to visualize both the axons of electroporated and non-electroporated RGCs. mRFP-, Lyn-SpiCee- and Lyn-SponGee-electroporated axons follow the TAG1-labeled tract, whereas SpiCee-Kras and SponGee-Kras expression led to axons defasciculating and growing at a distance of the chiasm before joining the rest of the retinal axons in the optic tract (Fig. 6b). This phenotype is reminiscent of the Slit1/2 double knock-out animals that also exhibit misguided axons at the optic chiasm^29,30^.

## Discussion

### Physiological relevance of second messenger compartmentation

Although a variety of second messenger microdomains have been described, the link between a biochemically-identified compartment and a function for such local signals has been scarcely studied. Investigating the physiological relevance of local second messenger signaling has remained challenging until recently with the lack of tools enabling to manipulate cAMP, cGMP and Ca^2+^ in subcellular domains of known biochemical identity. The development of genetically-encoded scavengers buffering cAMP, cGMP and Ca^2+^ has open a way to control the concentration of these signaling molecules with subcellular resolution^33,38,39^. Using this approach, we provide a comprehensive description of the subcellular compartmentation of cyclic nucleotides and Ca^2+^ signals induced in retinal growth cones by two repellent axon guidance molecules. Surprisingly, we found that ephrin-A5 and Slit1 downstream signaling involve second messenger signaling in distinct submembrane compartments: lipid rafts and the non-raft domain of the plasma membrane, respectively. The interactions between cAMP, cGMP and Ca^2+^ are also domain-specific. We confirmed that these differences in signaling correlate with distinct behaviors of axons exposed to either ephrin-A5 or Slit1 *in vitro. In vivo*, altering second messenger signaling in or outside lipid rafts induces distinct axon guidance phenotypes. Lipid raft-restricted signaling controls axon pathfinding in areas of the brain where ephrin-A5 is involved, whereas lipid raft-excluded second messenger modulation is required where Slit1 is critical for retinal axon guidance. A similar approach has been used to demonstrate that cAMP and cGMP signaling restricted to the primary cilium or to the closely apposed centrosome regulate distinct features of cortical interneuron migration (Atkins et al., co-submitted manuscript). Overall, these studies demonstrate that the restriction of second messengers in micro or nanodomains is functionally relevant and is critical for the development of the nervous system. The approach used is adaptable to other systems to enlarge the investigation of the physiological processes requiring second messenger signals restricted to subcellular compartments.

### Integration of multiple axon guidance cues occurs downstream of second messengers

The idea that second messenger signaling restricted to micro or nanodomains might activate distinct effectors downstream of axon guidance molecules has been formulated in particular to differentiate axon attractants and repellents^42^. However, the model accepted in most cases is that second messengers are placed in such a position in the signaling pathway of axon guidance molecules that they play the role of integrators. They might thus enable a switch between attraction and repulsion in a growth cone exposed to a mix of several guidance molecules. For instance, inverting the cAMP:cGMP ratio is sufficient to convert Netrin-1-induced attraction of spinal axons into repulsion^25^. The overall model to explain how Ca^2+^ can regulate both attractive and repulsive behaviors of the growth cone is based on codes relying on signal amplitude of the response and the slope of the calcium gradient across the growth cone^42^. Our observations rather demonstrate that different axon guidance molecules induce second messenger signals in distinct compartments, thus placing cAMP, cGMP and Ca^2+^ above a potential molecular integrator. Such an integrator should be in a position to combine second messenger signaling from different compartments of the growth cone.

Strikingly, the second messenger signals regulate the behavior of developing axons facing distinct axon repellents, thus modulating subtle changes in growing axons. Lipid raft-excluded second messenger variations regulate the response to Slit1, an axon guidance cue that controls the path of growing retinal axons still at a distance from their target^29,30^. This axon repellent thus does not terminate the growth of the axon and enable its further extension in the optic tract. This is in line with the observation that Slit1 does not induce an immediate retraction *in vitro*. By contrast, ephrin-A5 is involved in shaping retinal projections in areas of the brain where axons form their terminal arbors and thus require a stop signal preventing further extension^31,41^. This stop signal involves second messenger signaling in lipid rafts. Consistently, ephrin-A5 induces the retraction of the axons in the minutes following their bath application *in vitro*. These observations highlight that within the family of axon repellents, distinct axonal behaviors are regulated by cAMP, cGMP and Ca^2+^ signals confined in different subcellular compartments, thus placing these second messengers upstream of a final process that integrates the diversity of guidance stimuli present in the environment of developing axons.

### Expanding the repertoire of available second messenger codes

The wide range of signaling pathways regulated by cyclic nucleotides and Ca^2+^ requires a mechanism leading second messenger signals to achieve specificity for their downstream effector without interfering with the other cellular processes that these signaling molecules regulate. The subcellular compartmentation of cAMP, cGMP and Ca^2+^ contributes to such specificity. For instance, subcellular-restricted signals have been identified in the filopodia of retinal growth cones, in T-tubule of cardiomyocytes, in lipid rafts or in the primary cilium^5,9,10,43^. Here, we provide evidence that in addition to subcellular compartmentation of second messengers, the crosstalk between cAMP, cGMP and Ca^2+^ is highly controlled at a subcellular scale, with distinct interactions in the vicinity of lipid rafts and further away from this membrane compartment. Similarly cAMP and cGMP buffering in the primary cilium affects the polarity of migrating cortical interneurons in an opposite manner, whereas preventing the modulation of cyclic nucleotides at the centrosome leads to the same alteration of nucleokinesis, thus suggesting distinct co-regulations of these signaling molecules in these two compartments (Atkins et al., co-submitted manuscript). This is in line with the previously reported interactions between these signaling molecules, which are not conserved across cell types, suggesting multiple molecular controls of second messenger concentrations. Whereas Ca^2+^ drives the drop of cAMP concentration in insulin-secreting MIN6 β-cells exhibiting combined cAMP and Ca^2+^ oscillations, it is placed upstream of an elevation in cAMP in HEK cells with similar second messenger oscillations or in developing neurons^18–20^. However, these interactions were identified using cell-wide imaging and pharmacological manipulations, *i*.*e*. without subcellular resolution. Using such approaches, the detected crosstalk is thus likely to reflect the overall dominant pathway in each cell type, but might not report the diversity of second messenger interplays in distinct compartments within the same cell.

Achieving specificity by subcellular second messenger signals is limited by the number of available cellular domains that might not be sufficient to control the myriad of cAMP-, cGMP- and Ca^2+^-dependent signaling pathways. A diversity of crosstalks activated by distinct stimuli within the same cellular compartment might be a mechanism enabling improved specificity. Within a subcellular domain, it is conceivable that the combination of second messenger modulations and their crosstalk might differ downstream of distinct receptors or signaling pathways, thus expanding the range of available second messenger codes when merging the diversity of subcellular domains with the different signal combinations or crosstalks. This enlarged set of signals would enable to specifically activate a wider range of downstream effectors.

## Acknowledgements

We thank Dr O. Griesbeck for the gift of Twitch2b. We are grateful to Christine Métin, Melody Atkins, Camille Michaud and the members of our lab for thoughtful discussion and helpful critical reading of the manuscript, and to the members of the animal and imaging facilities of Institut de la Vision. This work was supported, by grants from ANR (ANR-18-CE16-0017) and Fondation pour la Recherche Médicale (EQU202003010158) to X.N. This work was performed in the frame of LABEX LIFESENSES (ANR-10-LABX-65) and of IHU FOReSIGHT (ANR-18-IAHU-0001) supported by French state funds managed by the Agence Nationale de la Recherche within the Investissements d’Avenir program. S.B. was supported by fellowships from the ED3C doctoral program (Sorbonne Université) and Fondation pour la Recherche Médicale (FDT202106013022). J.B. was supported by a fellowship from Fondation de France (00099274).

## Author contributions

Conceptualization, OR, XN; Methodology, SB, YZ, OR, XN; Validation, SB, OR, XN; Formal Analysis, SB, YZ, CGB, JB, OR, XN; Investigation, SB, YZ, FR, CGB, JB, MB, SC; Writing – Original Draft, SB, XN; Writing – Review & Editing, SB, YZ, FR, CGB, JB, MB, SC, OR, XN; Visualization, SB, XN; Supervision, OR, XN; Project Administration, XN; Funding acquisition, XN.

## Materials & Correspondence

The plasmids used in this study can be obtained from Addgene or from XN.

## Supplementary Material

Materials and Methods

Figs. S1 to S3

Tables S1 Movie S1

## Methods

### Animals

Pregnant C57BL6/J and RjOrl:SWISS mice were purchased from Janvier Labs. All animal procedures were performed in accordance with institutional guidelines and approved by the local ethics committee (C2EA-05: Comité d’éthique en expérimentation animale Charles Darwin). Animals were housed on a 12 h light/12 h dark cycle. Embryos from dated matings (developmental stage stated in each section describing individual experiments) were not sexed during the experiments and the female over male ratio is expected to be close to 1.

### Retinal explants

Retinas of E14.5 mouse embryos were electroporated with using two poring pulses (square wave, 175 V, 5 ms duration, with 50 ms interval) followed by four transfer pulses (40 V, 50 ms and 950 ms interpulse) with a Nepa21 Super Electroporator (NepaGene). This electroporation procedure was used for the following plasmids: mRFP, Lyn-SpiCee, mut-Lyn-SpiCee, SpiCee, SpiCee-Kras, mut-SpiCee-Kras, SponGee, Lyn-SponGee, mut-Lyn-SponGee, SponGee-Kras, mut-SponGee-Kras, cAMP Sponge, Lyn-Aspx, Aspx-Kras, mut-Aspx-kras, Lyn-H147, H147-Kras, Twitch2B, Lyn-Twitch2b, Twitch2b-Kras, ^T^HPDE5^VV^ (2 μg·μL^-1^ for single plasmid electroporation, 1 μg·μL^-1^ for each plasmid for co-electroporation). Retinas were dissected and kept 24 hours in culture medium (DMEM-F12 supplemented with 1 mM glutamine (Sigma Aldrich), 1% penicillin/streptomycin (Sigma Aldrich), 0.001% BSA (Sigma Aldrich) and 0.07% glucose), in a humidified incubator at 37°C and 5% CO2. The following day, the retinas were cut into 200 μm squares with a Tissue-Chopper (McIlwan) and explants were plated on glass coverslips or ibidi 35mm glass bottom dish coated with 100 μg·mL^-1^ poly-lysine and 20 μg·mL^-1^ Laminin (Sigma Aldrich). Cells were cultured for 24 hours in culture medium supplemented with 0.5% (w/v) methyl cellulose and B-27 (1/50, Life technologies).

### HEK293 cell culture

HEK293T cells (ATCC, not authenticated, free of mycoplasma contamination) were maintained in a 37°C, 5% CO2 incubator and transfected using Lipofectamine 2000 (Life Technologies) according to the manufacturer’s protocol and processed for membrane fractionation the day following transfection.

### Molecular Biology

Plasmids generated in this study (Lyn-Twitch2b, Twitch2b-Kras, Lyn-mut SponGee, mut SponGee-Kras, Lyn-mut SpiCee, mut SpiCee-Kras) are available upon request. The other plasmids used have been deposited to Addgene under the following name and catalog number: SpiCee, #140836; Lyn-SpiCee, #140837; SpiCee-Kras, #140838; SponGee, #134775; Lyn-SponGee, #134776; SponGee-Kras, #134777; pCX-^T^hPDE5^VV^, #134778.

Mut SponGee carries the E247G and E299G mutations in the cGMP binding sites. The Lyn-mut SponGee insert was generated by gene synthesis (Thermofisher) and was subcloned into an empty pCX vector (containing a CAG promoter) between the Acc65I and NotI restriction sites. The Lyn targeting sequence flanked by two NheI sites was removed from Lyn-mut SponGee using NheI. Mut SponGee was then inserted into a mRFP-Kras-containing pCX plasmid between the AgeI and BsrGI restriction sites.

Mut SpiCee carries the following mutations: D52A, E63Q, D91A, E102Q, D135A, E146Q, D171A and E182Q. Mut SpiCee was subcloned into a Lyn-SpiCee vector between the BbvCI and BsrGI restriction sites, thus replacing the SpiCee sequence and generating the Lyn-mut SpiCee construct carrying a CAG promoter. Mut SpiCee was also subcloned into a SpiCee-Kras vector between the AgeI and BsrGI restriction sites, thus replacing the SpiCee sequence and generating the mut SpiCee-Kras construct under the control of a CAG promoter.

A plasmid containing a CAG promoter followed by a Lyn targeting sequence in frame with the Twitch2b sequence, a stop codon and the Kras sequence was generated using the In-Fusion kit (Ozyme) and oligonucleotides for the Lyn and Kras sequences. The Lyn sequence and the stop codon preceding the Kras sequence were excised using AgeI and AflII, respectively, to obtain the Twitch2b-Kras construct, whereas the Kras sequence was removed using NheI to obtain the Lyn-Twitch2b-encoding plasmid.

All plasmids were sequenced to verify the success of the cloning strategies

### Membrane fractionation by detergent-free method

Electroporated retinas were pelleted (195 g for 5 min at 4°C) and resuspended in 1.34 mL of 0.5 M sodium carbonate, pH 11.5, with protease inhibitor cocktail and phosphatase inhibitor cocktails 1, 2 and 3 (Sigma-Aldrich). The homogenate was sheared through a 26-gauge needle and sonicated three times for 20-second bursts. The homogenate was adjusted to 40% (w/v) sucrose by adding 2.06 mL of 60% (w/v) sucrose in MBS (25 mM MES, pH 6.4, 150 mM NaCl, and 250 mM sodium carbonate), placed under a 5–30% (w/v) discontinuous sucrose gradient, and centrifuged at 34,000 rpm for 15–18 h at 4°C in a Beckman SW 41Ti rotor. Nine fractions (1.24 mL each) were harvested from the top of the tube mixed with 9 volumes of MBS, and centrifuged at 40,000 rpm for 1 h at 4°C (Beckman SW-41Ti rotor). Supernatants were discarded, and membrane pellets were resuspended in 100 μL of 1% (w/v) SDS.

For immunoblotting, samples were separated on a precast gel (4-15% Mini-Protean TGX Tris-Glycine-buffer SDS PAGE, Biorad) and transferred onto 0.2 μm Trans-Blot Turbo nitrocellulose membranes (Biorad). Membranes were blocked for one hour at room temperature in 1xTBS (10 mM Tris pH 8.0, 150 mM NaCl) supplemented with 5% (w/v) dried skim milk powder. Primary antibody incubation was carried out overnight at 4°C, with the following antibodies: rabbit anti-DsRed (1/200; 632476; Clontech; lot # 1306037), rabbit anti-β-Adaptin (1/200; sc-10762; Santa Cruz; lot # E1304) and rabbit anti-Caveolin (1/500; 610060; BD Transduction Laboratories; lot # GR256941-5). All primary antibodies have been previously validated for this assay ^9^. A goat anti-rabbit-HRP-coupled secondary antibody was used for detection (Jackson ImmunoResearch, West Grove, PA). After antibody incubations, membranes were extensively washed in TBS T (TBS containing 2.5% (v/v) Tween-20). Western blots were visualized using the enhanced chemiluminescence method (ECL prime Western Blotting detection reagent, Amersham).

### In utero retinal electroporation

*In utero* electroporation was performed like previously described ^44,45^. In brief, timed-pregnant mice (Janvier Labs) were delivered to the animal facility a week prior to the surgery in order to allow a minimum of 5 days adaptation. C57BL/6NRj pregnant mice were anesthetized with an intraperitoneal injection of a Xylazine/Ketamine mix (10 mg·kg^-1^ and 100 mg·kg^-1^, respectively) and a subcutaneous injection of buprenorphine (0.0125 mg·kg^-1^) was made pre-surgery for analgesia. Midline laparotomy was performed, exposing uterine horns and allowing visualization of embryos. The left eye of E14.5 embryos was injected with 2 μg·μL^-1^ of DNA using an elongated glass capillary (Harvard apparatus) with different plasmid solutions. The success of DNA injection was assessed using 0.07% fast green supplemented to the DNA solution. The eye was then electroporated with 5 pulses of 45 V during 50 ms every 950 ms (Nepagene electroporator). To target the central part of the retina, the positive electrode (CUY650P5, Sonidel) was placed on the side of the injected eye. Following surgery, the incision site was sutured (4-0, Ethicon), and mice were allowed to give birth. To increase the survival of the electroporated pups, a Swiss mouse was housed together with the mice that underwent surgery to favor the care of the pups. The Swiss mouse, mated a day earlier than the C57BL/6NRj mice, gave birth one day earlier. At P0, only 2 Swiss pups were left in the cage so that the electroporated pups were adopted by the Swiss mouse.

### Collapse assay

Retinal explants were treated with 200 ng·mL^-1^ rmSlit-1 or 500 ng·mL^-1^ rmEphrinA5 (R&D Systems) diluted in warm culture medium for 20 minutes before fixation with 4% (w/v) PFA in Sucrose 4% (w/v) for 30 minutes.

### Immunostaining following collapse assay

Retinal explants were permeabilized and blocked with 0.25% (v/v) Triton and 3% (w/v) BSA in PBS, then immunized against DsRed (1/1000, Takara Bio, lot #CDSO0219101) followed by a secondary antibody coupled to AlexaFluor 594 (1/500, Thermo Fisher Scientific) and βIII-tubulin (1//1000, Biolegend, lot #B249869) followed by a secondary antibody coupled to AlexaFluor 488 (1/500, Thermo Fisher Scientific). Antibodies were diluted in PBS supplemented with 0.1% (v/v) Triton and 1% (w/v) BSA.

### Whole mount immunostaining and tissue clearing

P15 mice were deeply anesthetized with a mix of Xylazine/Ketamine (20 mg.kg^-1^ and 200 mg.kg^-1^ respectively), perfused transcardially with 4% PFA in 0.12 M phosphate buffer. Retinas and brains were dissected out and post-fixed 12 hours in 4% PFA. Retinas (oriented with an incision on the ventral part) and were mounted in Mowiol. To validate area of electroporation, retinas were imaged under a 2.5X objective using an epi-fluorescence microscope (Leica DMI6000B). E18.5 embryos were harvested and the heads were fixed 3 hours with 4% PFA. The skulls were dissected out to harvest the brain and postfixed 12 hours in 4% PFA.

The samples were prepared according to the iDISCO+ protocol adapted from ^46^. The brains were then dehydrated in succeeding baths of methanol/PBS for 1.5 hours each at RT (50% MeOH, 80% MeOH, 100% MeOH). The samples were then transferred overnight in a depigmentation solution of methanol containing 6% H_2_O_2_ (VWR, 216763) at 4°C. E18.5 samples were then depigmented for 3 additional days in a solution of methanol containing 10% H_2_O_2_ at 4°C.

The samples were rehydrated in succeeding bath of methanol/PBS for 1.5 hours each at room temperature (100% MeOH X2, 80% MeOH, and 50% MeOH, PBS) and kept in PBS at 4°C before immunostaining.

The brains were permeabilized in the blocking solution (0.5% Triton-X100, 0.2% gelatin, 1XPBS, 0.1 g/L thimerosal) for 2 days for P15 brains and 24 hours for E18.5 brains at room temperature on agitation. For immunostaining, the samples were incubated with the primary antibodies against TAG1 (1/500, R&D Systems, lot #CDSO0219101) and or DsRed (1/1000, Takara Bio, lot #2103116) in a solution containing 0.5% Triton-X100, 0.2% gelatin, 1XPBS, 0.1g/L thimerosal, 10mg/mL saponin. Incubation in the primary antibody solutions was performed for 2 weeks for P15 brains or 1 week for E18.5 brains at 37°C under agitation. The samples were washed for 1 day. The samples were then immunized by secondary antibodies coupled to AlexaFluor 647 (1/500, Jackson Immunoresearch, lot #146920) and or Cy3 (1/500, Jackson Immunoresearch, lot #148687) diluted in the same solution as for the primary antibodies, passed through a 0.22 μm filter and incubated for 1 week for P15 brains and 2 days for E18.5 brains at 37°C under agitation. The samples were then washed for 6 times during 1 day in PBS supplemented with 0.2% gelatin and 0.5% Triton-X100, and 2 washes of 1X PBS prior to storing the samples in the dark at 4°C until clearing.

For clearing, the samples were first transferred overnight in a solution containing 20% MeOH. The following day they were incubated in succeeding bath of Methanol/PBS for 1 hour (40% MeOH, 60% MeOH, 80% MeOH, 100% MeOH X2). Sample were then placed over night in a solution of 2/3 dichloromethane and 1/3 MeOH. The following day, they were incubated 30 min in a 100% dichloromethane bath prior being transferred in 100% benzyl ether (DBE) and stored until imaging.

### Light sheet microscopy

Images were acquired with an ultramicroscope I (LaVision BioTec, Miltenyi Biotec) coupled to a 2x objective (Olympus, MVPLAPO) with different magnifications (0.63x, 1x, 1.25x, 1.6x, 2x, 2.5x, 3.2x, 4x and 5x) or a 12x objective and with the ImspectorPro software (LaVision Biotec, Miltenyi Biotec). The light sheet was generated by a laser (wavelength 561 and 640 nm, coherent Sapphire Laser, LaVision BioTec, Miltenyi Biotec). Samples were imaged in DBE with a Zyla SCMOS camera (Andor, Oxford Instrument). Step size between each image was fixed at 1 μm (NA = 0.5, 150 ms time exposure).

### FRET Imaging and analysis

Images were acquired with an inverted DMI6000B epifluorescence microscope (Leica) coupled to a 40x oil-immersion objective (N.A. 1.3) and Metamorph software (Molecular Devices). Retinal explants were perfused (0.3 mL min -1) with 1 mM CaCl2, 0.3 mM MgCl2, 0.5 mM Na2HPO4, 0.45 mM NaH 2PO4, 0.4 mM MgSO4, 4.25 mM KCl, 14 mM NaHCO3, 120 mM NaCl, 0.0004% CuSO4, 0.124 mM Fe(NO3)3, 1.5 mM FeSO4, 1.5 mM thymidine, 0.51 mM lipoic acid, 1.5 mM ZnSO4, 0.5 mM sodium pyruvate (all from Sigma), 1X MEM Amino Acids (Life Technologies), 1X non-essential amino acids (Life Technologies), 25 mM HEPES (Sigma), 0.5 mM putrescine (Sigma), 0.01% BSA (Sigma), 0.46% glucose (Sigma), 1 mM glutamine (Life Technologies), 2% penicillin streptomycin (Life Technologies). Vitamin B12 and riboflavin were omitted because of their auto-fluorescence. rmSlit-1 was used at 200 ng·mL^-1^ and rmEphrinA5 at 500 ng·mL^-1^ (R&D Systems). Spermine-NONOate was used at 50mM, Forskolin (Sigma) at 10mM and ionomycin (Invitrogen) was used at 5 μM. Images were acquired simultaneously for the CFP (483/32 nm) and YFP (542/27) channels every 20 seconds for cAMP or cGMP detection, or every 5 seconds for calcium detection, while cells were continuously superfused with the medium described above. Simultaneous CFP and YFP channel acquisition was achieved using a dual chip CCD camera ORCA-D2 (Hamamatsu). The wavelength used for CFP excitation was 436/20 nm. Images were processed in ImageJ, corrected for background and bleedthrough from CFP into the YFP channel, and the CFP:YFP (H147) or the YFP:CFP (^T^hPDE5^VV^, Twitch2b) ratio was computed and normalized to the initial values for each single cell.

### DIC Imaging and analysis

Images were acquired with an inverted Eclipse Ti2 (Nikon) coupled to a 40x oil-immersion objective and Metamorph software (Molecular Devices). Retinal explants were imaged in culture medium without phenol red and supplemented with 2 μM HEPES at 37°C. rmSlit-1 was used at 200 ng·mL^-1^ and rmEphrinA5 at 500 ng·mL^-1^ (R&D Systems). Images were acquired in DIC every 15 sec by using a Prime95B camera (Photometrics). Images were processed in ImageJ, growth length was measured with the Manual tracking plugin.

### Quantification and statistical analysis

No data were excluded from the analysis, except for FRET imaging for which axons lacking a NO-ionomycin- or Forskolin-induced change in FRET reflecting an elevation of cGMP, Ca^2+^ or cGMP, respectively, were excluded from the analysis. No sample size calculation was performed. Sample size was considered sufficient after at least three independent experiments, leading to n ≥ 3 since several animals, coverslips, or biochemical assays were often analyzed for the same experimental condition. Animals or cultures were equivalent and not distinguishable before treatment, *de facto* randomizing the sample without the need of a formal randomization process. Photomicrographs were often easily traceable by eye to its experimental condition, making blind analysis of the data difficult to achieve. When careful blinding was performed, experiments reproduced the results obtained in non-blinded experiments with identical experimental conditions. Image calculation and analysis were performed using ImageJ.

Statistical tests were calculated using GraphPad Prism (GraphPad Software Inc.). Supplementary Table 1 summarizes the number of replicates for all the data shown in Figures 1-7 and Supplementary Figures 1-3.

