## Supplemental Figure 1-3, Supplemental Table 1 and Supplemental Video legend for "Subcellular second messenger networks drive distinct repellent-induced axon behaviors"

**SUPPLEMENTARY FIGURES**

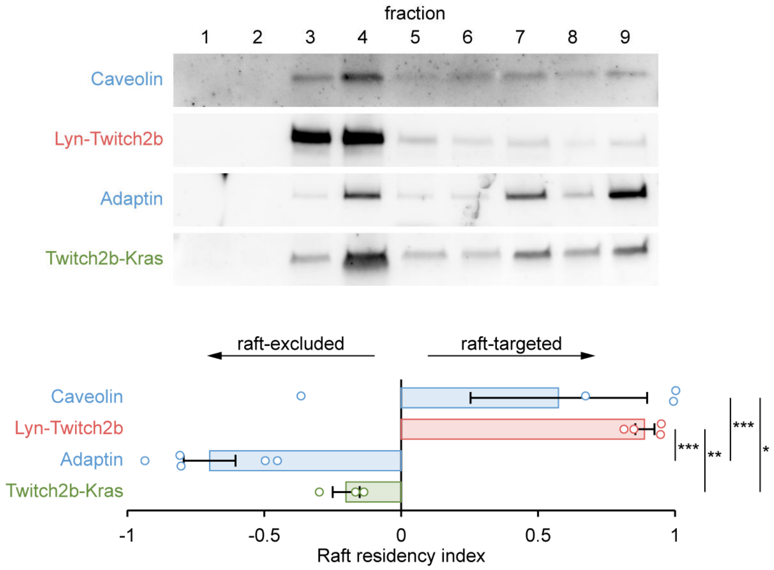


**Supplementary Figure 1. Subcellular targeting of the $Ca^{2+}$ biosensor Twitch2b.**
The FRET $Ca^{2+}$ biosensor Twitch2b was fused to a raft targeting- (Lyn) or a raft excluding-sequence (Kras) and electroporated in the developing retina. Membrane fractionation was performed to evaluate the subcellular localization of each construct. Lyn-Twitch2b was found in the same fraction as the raft-targeted protein Caveolin 1, whereas the pattern of Twitch2b-Kras mimics the distribution of Adaptin, a raft-excluded marker. The residency to lipid raft was quantified using a raft-targeting index calculated as ($I_{fraction\ 3}$ - $I_{fraction\ 9}$)/( $I_{fraction\ 3}$ + $I_{fraction\ 9}$), where $I_{fraction\ 3}$ and $I_{fraction\ 9}$ are the fraction of the signal found in fraction 3 and 9, respectively. Thus, this index ranges from -1 (lipid raft exclusion) to 1 (lipid raft targeting). The average, s.e.m. and individual data points are shown. * $P < 0.05$; ** $P < 0.01$; *** $P < 0.001$; One-way ANOVA.

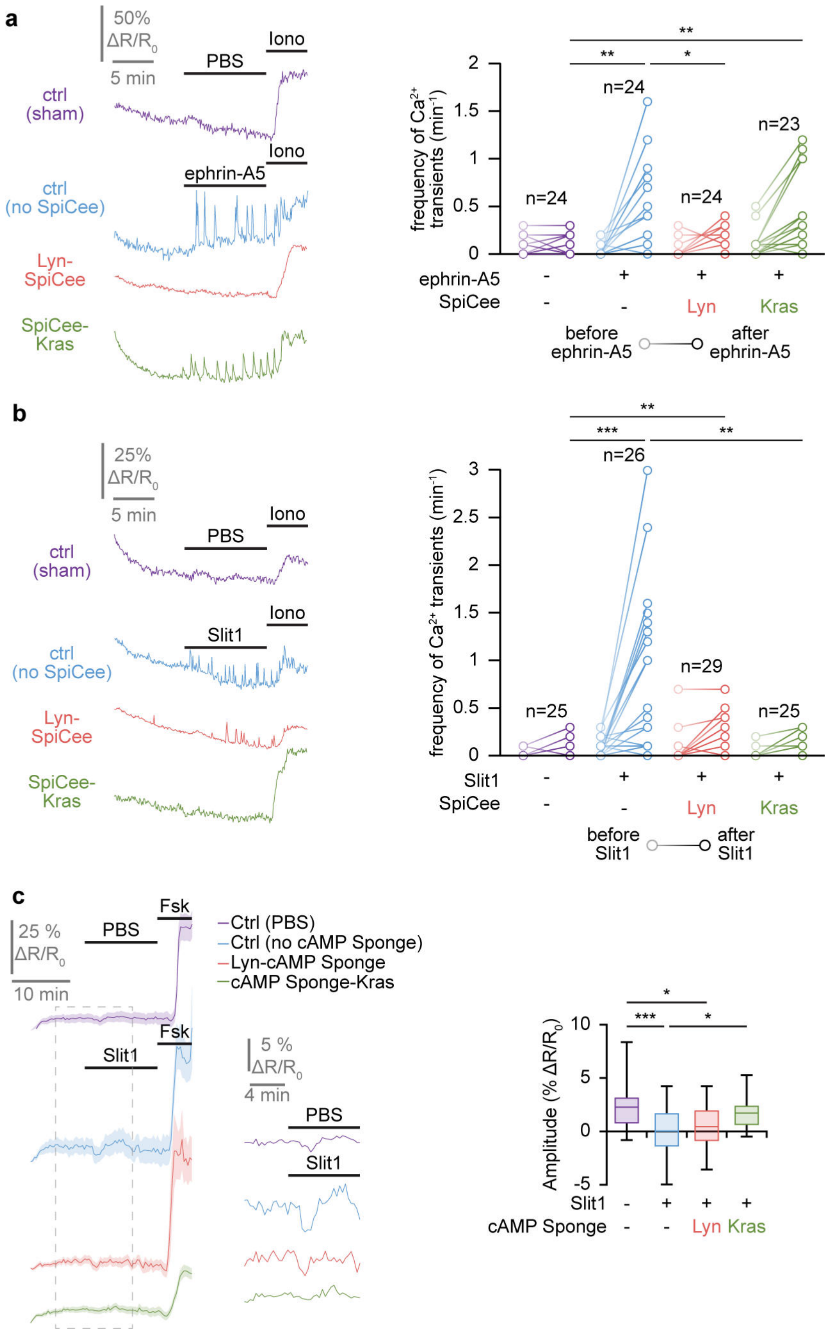


**Supplementary Figure 2. Specificity of the subcellular buffering of second messengers.**
**(a)** The lipid raft-targeted $Ca^{2+}$ biosensor Lyn-Twitch2b was expressed in retinal explants alone or in combination with the $Ca^{2+}$ scavengers targeted to (Lyn-SpiCee) or excluded from lipid rafts (SpiCee-Kras) and retinal axons were exposed to ephrin-A5. The ephrin-A5-induced and lipid raft-restricted elevation of $Ca^{2+}$ transient frequency (blue trace) was abolished by the lipid raft-restricted $Ca^{2+}$ scavenger Lyn-SpiCee (red trace), but not by its lipid raft-excluded equivalent SpiCee-Kras (green trace). A ionomycin (iono) stimulation leading to a $Ca^{2+}$ elevation is achieved at the end of each recording to verify the functionality of the biosensor and the viability of the axon. Representative traces are shown. Individual data points are shown. * $P < 0.05$; ** $P < 0.01$; Kruskal-Wallis test followed by Mann-Whitney post hoc tests. The number of quantified axons is indicated on the graphs.

**(b)** The lipid raft-excluded $Ca^{2+}$ biosensor Twitch2b-Kras was expressed in retinal explants alone or in combination with the $Ca^{2+}$ scavengers targeted to (Lyn-SpiCee) or excluded from lipid rafts (SpiCee-Kras) and retinal axons were exposed to Slit1. The Slit1-induced and lipid raft-excluded elevation of $Ca^{2+}$ transient frequency (blue trace) was abolished by the lipid raft-excluded $Ca^{2+}$ scavenger SpiCee-Kras (green trace), but not by its lipid raft-targeted equivalent Lyn-SpiCee (red trace). A ionomycin (iono) stimulation leading to a $Ca^{2+}$ elevation is achieved at the end of each recording to verify the functionality of the biosensor and the viability of the axon. Representative traces are shown. Individual data points are shown. ** $P < 0.01$; *** $P < 0.001$; Kruskal-Wallis test followed by Mann-Whitney post hoc tests. The number of quantified axons is indicated on the graphs.
**(c)** The lipid raft-excluded cAMP biosensor H147-Kras was expressed in retinal explants alone or in combination with the cAMP scavengers targeted to (Lyn-cAMP Sponge) or excluded from lipid rafts (cAMP Sponge-Kras) and retinal axons were exposed to Slit1. The Slit1-induced and lipid raft-excluded reduction in cAMP concentration (blue trace) was abolished by the lipid raft-excluded cAMP buffer cAMP Sponge-Kras (green trace), but not by its lipid raft-targeted equivalent Lyn-cAMP Sponge (red trace). A forskolin (Fsk) stimulation leading to a cAMP elevation is achieved at the end of each recording to verify the functionality of the biosensor and the viability of the axon. The portion of the left traces enclosed in the dashed rectangle is shown magnified in the right part of the panel. Shadows surrounding traces, s.e.m. Box-and-whisker plot elements: center line, mean; box limits, upper and lower quartiles; whiskers, $10^{th}$ and $90^{th}$ percentiles. * $P < 0.05$; *** $P < 0.001$; Kruskal-Wallis test followed by Mann-Whitney post hoc tests.

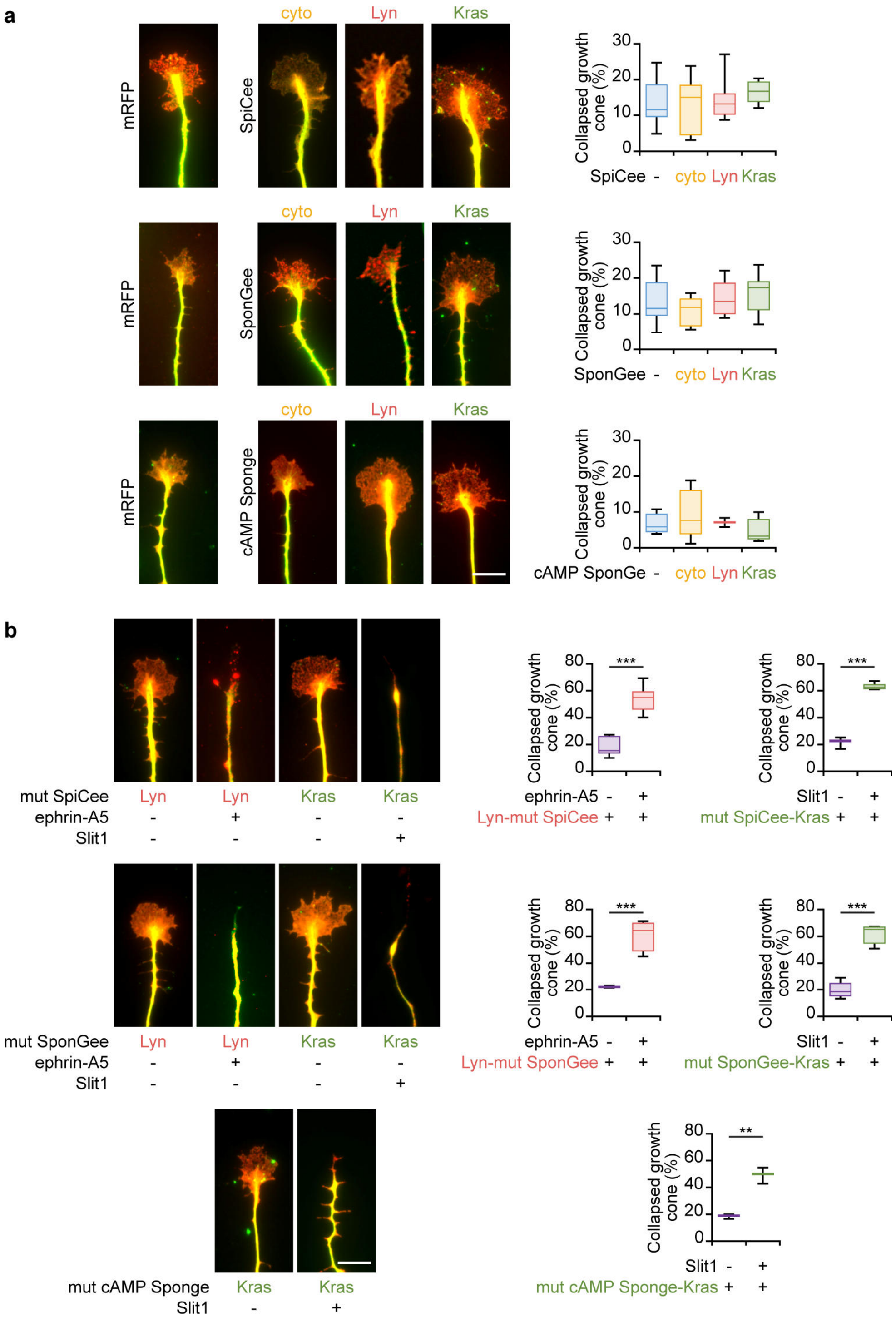


**Supplementary Figure 3. The morphology of growing axons is not affected by SpiCee, SponGee or cAMP Sponge expression and the second messenger binding sites of these scavengers are required for them to alter the collapse of RGC growth cones.**

**(a)** Retinal axons expressing SpiCee, SponGee or cAMP Sponge either lacking a targeting sequence (cyto), targeted to lipid rafts (Lyn) or excluded from this subcellular compartment (Kras) exhibit a morphology similar to mRFP-expressing axons. In particular, the number of collapsed growth is similar in all these experimental conditions. Scale bar, 10 µm. Box-and-whisker plot elements: center line, mean; box limits, upper and lower quartiles; whiskers, $10^{th}$ and $90^{th}$ percentiles. No statistically significant difference was found; one-way ANOVA.

**(b)** SpiCee, SponGee and cAMP Sponge carrying point mutations that abolish their ability to bind their targeted second messenger (mut SpiCee, mut SponGee and mut cAMP Sponge, respectively) do not affect the ephrin-A5- or Slit1-induced collapse of retinal growth cones. Axons were immunolabeled with a βIII-tubulin (green) and a Ds-Red (red) antibody. The latter reports the expression of SponGee, SpiCee, cAMP Sponge, or their targeted or mutated variants. Scale bar, 10 µm. Box-and-whisker plot elements: center line, mean; box limits, upper and lower quartiles; whiskers, $10^{th}$ and $90^{th}$ percentiles. ** $P < 0.01$; *** $P < 0.001$; Student t test.

### SUPPLEMENTARY TABLE

| Figure panel | Number of replicates |
|---|---|
| **Fig. 1b** | PBS, n=33 axons from 15 coverslips and 4 independent experiments. EphrinA5, n=22 axons from 14 coverslips and 8 independent experiments. SponGee, n=11 axons from 8 coverslips and 5 independent experiments. Lyn-SponGee, n=15 axons from 10 coverslips and 5 independent experiments. SponGee-Kras, n= 18 axons from 7 coverslips and 4 independent experiments. |
| **Fig. 1d** | Twitch2b PBS, n=20 axons from 9 coverslips and 3 independent experiments. Twitch2b ephrinA5, n=30 axons from 13 coverslips and 7 independent experiments. Lyn-Twitch2b PBS, n=24 axons from 6 coverslips and 3 independent experiments. Lyn-Twitch2b ephrinA5, n=24 axons from 7 coverslips and 5 independent experiments. Twitch2b-Kras PBS, n=25 axons from 8 coverslips and 3 independent experiments. Twitch2b-Kras ephrinA5, n=29 axons from 13 coverslips and 6 independent experiments. |
| **Fig. 2a** | H147 PBS, n=42 axons from 13 coverslips and 7 independent experiments. H147 slit1, n=42 axons from 9 coverslips and 7 independent experiments. Lyn-H147 PBS, n=22 axons from 8 coverslips and 6 independent experiments. Lyn-H147 slit1, n=35 axons from 12 coverslips and 6 independent experiments. H147-Kras PBS, n=40 axons from 12 coverslips and 6 independent experiments. H147-Kras slit1, n=34 axons from 9 coverslips and 6 independent experiments. |
| **Fig. 2b** | PBS, n=33 axons from 15 coverslips and 4 independent experiments. Slit1, n=18 axons from 9 coverslips and 3 independent experiments. SponGee, n=29 axons from 15 coverslips and 7 independent experiments. Lyn-SponGee, n=19 axons from 8 coverslips and 3 independent experiments. SponGee-Kras, n= 19 axons from 10 coverslips and 4 independent experiments. |
| **Fig. 2c** | Twitch2b PBS, n=20 axons from 9 coverslips and 3 independent experiments. Twitch2b slit1, n=25 axons from 11 coverslips and 3 independent experiments. Lyn-Twitch2b PBS, n=20 axons from 6 coverslips and 3 independent experiments. Lyn-Twitch2b slit1, n=29 axons from 9 coverslips and 3 independent experiments. Twitch2b-Kras PBS, n=25 axons from 8 coverslips and 3 independent experiments. Twitch2b-Kras slit1, n=26 axons from 15 coverslips and 6 independent experiments. |
| **Fig. 3a** | EphrinA5, n=22 axons from 14 coverslips and 8 independent experiments. Lyn-cAMP Sponge, n=31 axons from 5 coverslips and 3 independent experiments. Lyn-SpiCee, n=18 axons from 4 coverslips and 3 independent experiments. |
| **Fig. 3b** | Lyn-Twitch2b ephrin, n=24 axons from 7 coverslips and 5 independent experiments. Lyn-Twitch2b+Lyn-cAMP Sponge, n=29 axons from 8 coverslips and 4 independent experiments. Lyn-Twitch2b+Lyn-SponGee, n=33 axons from 10 coverslips and 3 independent experiments. |
| **Fig. 3c** | Lyn-H147, n=36 axons from 10 coverslips and 5 independent experiments. Lyn-H147+Lyn-SponGee, n=44 axons from 5 coverslips and 3 independent experiments. Lyn-H147+Lyn-SpiCee, n=20 axons from 7 coverslips and 3 independent experiments. |
| **Fig. 3e** | Slit1, n=18 axons from 9 coverslips and 3 independent experiments. cAMP Sponge-Kras, n=47 axons from 13 coverslips and 4 independent experiments. SpiCee-Kras, n=47 axons from 11 coverslips and 3 independent experiments. |

| | |
|---|---|
| **Fig. 3f** | Twitch2b-Kras, n=26 axons from 15 coverslips and 6 independent experiments. Twitch2b-Kras+cAMP Sponge-Kras, n=36 axons from 10 coverslips and 4 independent experiments. Twitch2b-Kras+SponGee-Kras, n=36 axons from 14 coverslips and 4 independent experiments. |
| **Fig. 3g** | H147-Kras, n=34 axons from 9 coverslips and 6 independent experiments. H147-Kras+SponGee-Kras, n=35 axons from 10 coverslips and 3 independent experiments. H147-Kras+SpiCee-Kras, n=26 axons from 4 coverslips and 3 independent experiments. |
| **Fig. 4a** | PBS, n=2095 axons from 18 coverslips and 8 independent experiments. EphrinA5, n=2269 axons from 11 coverslips and 5 independent experiments. SponGee, n=503 axons from 6 coverslips and 3 independent experiments. Lyn-SponGee, n=1226 axons from 6 coverslips and 3 independent experiments. SponGee-Kras, n= 862 axons from 6 coverslips and 3 independent experiments. |
| **Fig. 4b** | PBS, n=2095 axons from 18 coverslips and 8 independent experiments. EphrinA5, n=2269 axons from 11 coverslips and 5 independent experiments. SpiCee, n=1342 axons from 6 coverslips and 3 independent experiments. Lyn-SpiCee, n=962 axons from 6 coverslips and 3 independent experiments. SpiCee-Kras, n= 1306 axons from 6 coverslips and 3 independent experiments. |
| **Fig. 4c** | PBS, n=2095 axons from 18 coverslips and 8 independent experiments. Slit1, n=3450 axons from 22 coverslips and 8 independent experiments. SponGee, n=954 axons from 10 coverslips and 3 independent experiments. Lyn-SponGee, n=1570 axons from 9 coverslips and 3 independent experiments. SponGee-Kras, n=2471 axons from 16 coverslips and 4 independent experiments. |
| **Fig. 4d** | PBS, n=2095 axons from 18 coverslips and 8 independent experiments. Slit1, n=3450 axons from 22 coverslips and 8 independent experiments. SpiCee, n=1309 axons from 8 coverslips and 3 independent experiments. Lyn-SpiCee, n=1823 axons from 12 coverslips and 4 independent experiments. SpiCee-Kras, n=1397 axons from 8 coverslips and 3 independent experiments. |
| **Fig. 4e** | PBS, n=929 axons from 7 coverslips and 4 independent experiments. Slit1, n=948 axons from 8 coverslips and 4 independent experiments. cAMP Sponge, n=833 axons from 7 coverslips and 4 independent experiments. Lyn-cAMP Sponge, n=565 axons from 6 coverslips and 4 independent experiments. cAMP Sponge-Kras, n=334 axons from 4 coverslips and 3 independent experiments. |
| **Fig. 5** | PBS, n=70 axons from 15 coverslips and 6 independent experiments. Slit1, n=77 axons from 10 coverslips and 5 independent experiments. EphrinA5, n=57 axons from 8 coverslips and 5 independent experiments. |
| **Fig. 6a** | mRFP, n=9 brains from 4 independent experiments. Lyn-SpiCee, n=11 brains from 6 independent experiments. Lyn-SponGee, n=10 brains from 8 independent experiments. SpiCee-Kras, n=9 brains from 5independent experiments. SponGee-Kras, n= 8 brains from 4 independent experiments. |
| **Fig. 6b** | mRFP, n=16 brains from 7 independent experiments. Lyn-SpiCee, n=16 brains from 4 independent experiments. Lyn-SponGee, n=11 brains from 5 independent experiments. SpiCee-Kras, n=15 brains from 6 independent experiments. SponGee-Kras, n=11 brains from 5 independent experiments |
| **Supplementary Fig. 1** | Caveolin, n=4 independent experiments. Adaptin, n=5 independent experiments. Lyn-Twitch2b, n=4 independent experiments. Twitch2b-Kras, n=3 independent experiments. |

| | |
|---|---|
| **Supplementary Fig. 2a** | Lyn-Twitch2b PBS, n=24 axons from 5 coverslips and 3 independent experiments. Lyn-Twitch2b ephrinA5, n=24 axons from 7 coverslips and 5 independent experiments. Lyn-Twitch2b+Lyn-SpiCee, n=24 axons from 5 coverslips and 4 independent experiments. Lyn-Twitch2b+SpiCee-Kras, n=23 axons from 5 coverslips and 3 independent experiments. |
| **Supplementary Fig. 2b** | Twitch2b-Kras PBS, n=25 axons from 8 coverslips and 3 independent experiments. Twitch2b-Kras slit1, n=26 axons from 20 coverslips and 6 independent experiments. Twitch2b-Kras+Lyn-SpiCee, n=29 axons from 13 coverslips and 5 independent experiments. Twitch2b-Kras+SpiCee-Kras, n=25 axons from 4 coverslips and 3 independent experiments. |
| **Supplementary Fig. 2c** | H147-Kras PBS, n=40 axons from 12 coverslips and 6 independent experiments. H147-Kras slit1, n=34 axons from 8 coverslips and 5 independent experiments. H147-Kras+Lyn-cAMP Sponge n=26 axons from 10 coverslips and 5 independent experiments. H147-Kras+cAMP Sponge-Kras, n=45 axons from 11 coverslips and 4 independent experiments. |
| **Supplementary Fig. 3a** | SpiCee and SponGee dataset: mRFP, n=2095 axons from 18 coverslips and 8 independent experiments. SpiCee, n=721 axons from 7 coverslips and 3 independent experiments. Lyn-SpiCee n=1260 axons from 8 coverslips and 4 independent experiments. SpiCee-Kras, n=898 axons from 7 coverslips and 3 independent experiments. SponGee, n=636 axons from 5 coverslips and 3 independent experiments. Lyn-SponGee, n=1119 axons from 8 coverslips and 3 independent experiments. SponGee-Kras, n=1400 axons from 10 coverslips and 4 independent experiments.<br>cAMP Sponge dataset: mRFP, n=929 axons from 7 coverslips and 4 independent experiments. cAMP Sponge, n=766 axons from 6 coverslips and 3 independent experiments. Lyn-cAMP Sponge n=643 axons from 5 coverslips and 4 independent experiments. cAMP Sponge-Kras, n=416 axons from 4 coverslips and 3 independent experiments. |
| **Supplementary Fig. 3b** | Lyn-mut SpiCee PBS, n=1532 axons from 7 coverslips and 3 independent experiments. Lyn-mut SpiCee ephrinA5, n=2261 axons from 9 coverslips and 3 independent experiments. Lyn-mut SponGee PBS n=837 axons from 3 coverslips and 3 independent experiments. Lyn-mut SponGee ephrinA5, n=1370 axons from 6 coverslips and 3 independent experiments. mut SpiCee-Kras PBS, n=557 axons from 3 coverslips and 3 independent experiments. mut SpiCee-Kras slit1, n=863 axons from 3 coverslips and 3 independent experiments. mut SponGee-Kras PBS, n=1032 axons from 6 coverslips and 3 independent experiments. mut SponGee-Kras slit1, n=1215 axons from 9 coverslips and 3 independent experiments. mut cAMP Sponge-Kras PBS, n=380 axons from 3 coverslips and 3 independent experiments. mut cAMP Sponge-Kras slit1, n=321 axons from 3 coverslips and 3 independent experiments. |

**Supplementary Table 1. Summary of the number of replicates.**

### SUPPLEMENTARY MOVIE LEGEND

**Supplementary Movie 1. Ephrin-A5 and Slit1 induce distinct morphological changes of axonal growth cones *in vitro*.**

The growth of axons exposed to PBS was not affected (left video). Ephrin-A5 induced a growth cone collapse followed by a prompt retraction (middle video). Axons exposed to Slit1 exhibited a collapse of the growth cones but in contrast to axons encountering ephrin-A5, do not retract within the 20 minutes recorded (right video).
